# Genomic landscape of the global oak phylogeny

**DOI:** 10.1101/587253

**Authors:** Andrew L. Hipp, Paul S. Manos, Marlene Hahn, Michael Avishai, Cathérine Bodénès, Jeannine Cavender-Bares, Andrew Crowl, Min Deng, Thomas Denk, Sorel Fitz-Gibbon, Oliver Gailing, M. Socorro González-Elizondo, Antonio González-Rodríguez, Guido W. Grimm, Xiao-Long Jiang, Antoine Kremer, Isabelle Lesur, John D. McVay, Christophe Plomion, Hernando Rodríguez-Correa, Ernst-Detlef Schulze, Marco C. Simeone, Victoria L. Sork, Susana Valencia-Avalos

## Abstract

- The tree of life is highly reticulate, with the history of population divergence buried amongst phylogenies deriving from introgression and lineage sorting. In this study, we test the hypothesis that there are regions of the oak (*Quercus*, Fagaceae) genome that are broadly informative about phylogeny and investigate global patterns of oak diversity.
- We utilize fossil data and restriction-site associated DNA sequencing (RAD-seq) for 632 individuals representing ca. 250 oak species to infer a time-calibrated phylogeny of the world’s oaks. We use reversible-jump MCMC to reconstruct shifts in lineage diversification rates, accounting for among-clade sampling biases. We then map the > 20,000 RAD-seq loci back to a recently published oak genome and investigate genomic distribution of introgression and phylogenetic support across the phylogeny.
- Oak lineages have diversified among geographic regions, followed by ecological divergence within regions, in the Americas and Eurasia. Roughly 60% of oak diversity traces back to four clades that experienced increases in net diversification due to climatic transitions or ecological opportunity.
- The support we find for the phylogeny contrasts with high genomic heterogeneity in phylogenetic signal and introgression. Oaks are phylogenomic mosaics, and their diversity may in fact depend on the gene flow that shapes the oak genome.

## Introduction

The tree of life exhibits reticulation from its base to its tips (Folk *et al*., 2018; Quammen, 2018). Oaks (*Quercus* L., Fagaceae) are no exception (Hipp, 2018), and in fact the genus is rife with case-studies in localized gene flow (e.g. Hardin, 1975; Whittemore & Schaal, 1991; McVay *et al*., 2017a; Kim *et al*., 2018), and ancient introgression (Crowl *et al*., In review; McVay *et al*., 2017b; Kim *et al*., 2018). Oaks have in fact been held up as a paradigmatic syngameon (Hardin, 1975; Van Valen, 1976; Dodd & Afzal-Rafii, 2004; Cannon & Scher, 2017; Boecklen, 2017), a system of interbreeding species in which incomplete reproductive isolation may facilitate adaptive gene flow and species migration (Petit *et al*., 2003; Dodd & Afzal-Rafii, 2004). The oak genome (Plomion *et al*., 2018) consequently tracks numerous unique species-level phylogenetic histories that result from lineage sorting and differential rates of introgression (Anderson, 1953; Eaton *et al*., 2015; McVay *et al*., 2017b; Edelman *et al*., 2018). Oak genomes are mosaics of disparate phylogenetic histories (cf. Pääbo, 2003). Given the prevalence of hybridization in trees globally (Petit & Hampe, 2006; Cannon & Lerdau, 2015), understanding how these stories line up with one another, and whether there are regions of the genome that track a common story, is essential to understanding the prevalence of adaptive gene flow and the phylogenetic history of forest trees.

Restriction-site associated DNA sequencing (RAD-seq; Miller *et al*., 2007a,b; Lewis *et al*., 2007; Baird *et al*., 2008; Ree & Hipp, 2015) has revolutionized our understanding of oak phylogeny in the past five years (Jiang *et al*., In review; Hipp *et al*., 2014, 2018; Cavender-Bares *et al*., 2015; Eaton *et al*., 2015; Hipp, 2017; Fitz-Gibbon *et al*., 2017; Pham *et al*., 2017; Ortego *et al*., 2018; Deng *et al*., 2018; Kim *et al*., 2018). Its ties to the genome, however, have not been fully exploited because of the lack of an assembled genome. While earlier studies have explored the effects of gene identity on phylogenetic informativeness (Hipp *et al*., 2014) and genomic heterogeneity in phylogenetic vs. introgressive signals (McVay *et al*., 2017b,a), they have not had access to the oak genome sequence. As a consequence, we do not understand the distribution of genomic breakpoints between introgressive and divergent histories. Moreover, no studies to date have brought together a comprehensive sampling of taxa to investigate the history of diversification across the genus.

In this paper, we integrate data from the recently published *Quercus robur* genome (Plomion *et al*., 2016, 2018) with previously published RAD-seq data for 427 sequenced oak individuals across the tree of life and new RAD-seq data for an additional 205 individuals to investigate the global oak phylogenomic mosaic for approximately 60% of the world’s *Quercus* species. We test the hypothesis that there are regions of the genome that are uniformly informative about *Quercus* phylogeny, regions that make oak lineages what they are. Furthermore, using a time-calibrated one-tip-per species tree novel to this study for ca. 60% of known species, we test the hypothesis that the high diversity of oaks in Mexico and eastern China is a consequence of high diversification rates. Finally, we show that the consensus of the evolutionary histories of more than 20,000 RAD-seq loci matches our understanding of oak evolution based on morphological information from extant and fossil species in spite of broadly conflicting individual locus genealogies.

## Materials and Methods

### Previously published RAD-seq and new RAD-seq: sequencing and clustering

Data from previously published RAD-seq phylogenies were analyzed alongside new RAD-seq data for a total of 632 individuals (Table S1). RAD-seq data were generated as described in the previous studies. New data were from library preparations conducted at Floragenex, Inc. (Portland, OR, USA) following the methods of Baird *et al*. (2008) with *Pst*I, barcoded by individual, and sequenced on an Illumina Genome Analyzer IIx at Floragenex, or an Illumina HiSeq 2500 or HiSeq 4000 at the University of Oregon Genomic Facility.

FASTQ files were demultiplexed and filtered to remove sequences with more than 5 bases of quality score < 20 and assembled into loci for phylogenetic analysis using ipyrad 0.7.23 (Eaton, 2014) at 85% sequence similarity. Consensus sequences for each individual for each locus were then clustered across individuals, retaining loci present in at least 4 individuals and possessing a maximum of 20 SNPs and 8 indels across individuals. The dataset was filtered to loci with a minimum of 15 individuals each, for a total of 58,985 loci. Data were imported into R using the RADami package (Hipp *et al*., 2014) for downstream analysis.

RAD-seq loci were mapped back to the latest version of the *Quercus robur* haploid genome (haplome 2.3; https://urgi.versailles.inra.fr/Data/Genome/Genome-data-access) (Plomion *et al*., 2018). The oak genome is made of 12 pseudomolecules (*i.e*. chromosomes) and a set of 538 unassigned scaffolds. Mapping was performed using Blast+ 2.8.1 (Camacho *et al*., 2009). We filtered alignments based on expect (E) values (E-value ≤10^−5^), alignment length (≥80% of the length of the loci) and percent identity (≥80%). For each locus, the best alignment was kept. All sequence data analyzed in this paper are available as FASTQ files from NCBI’s Short Read Archive (Table S1), and aligned loci and additional data and scripts for all analysis are available from https://github.com/andrew-hipp/global-oaks-2019. Analysis details are in the Supplement (Methods S1).

### Phylogenetic analysis

Maximum likelihood phylogenetic analyses were conducted in RAxML v8.2.4 (Stamatakis, 2014) using the GTRCAT implementation of the general time reversible model of nucleotide evolution (Stamatakis, 2006), with branch support assessed using RELL bootstrapping (Minh *et al*., 2013). For the phylogeny including all tips (Fig. S1), analysis was unconstrained, and we used the taxonomic disparity index (TDI) of Pham *et al*. (2016) to identify the extent of non-monophyly by species. Topology within the white oaks of sections *Ponticae*, *Virentes*, and *Quercus* (hereafter in the paper “white oaks *s.l.*,” contrasted with “white oaks *s.str.*” for just section *Quercus*) was observed to be at odds with previous close studies (Crowl *et al*., In review; McVay *et al*., 2017b,a; Hipp *et al*., 2018) that have shown the topology of the white oaks *s.l.* to be sensitive to taxon and locus sampling. For dating, samples were pruned to one sample per named species, favoring samples with the most loci, except for species in which variable position of samples from different populations was deemed to represent cryptic diversity, in which case more than one exemplar was retained. The singletons tree was estimated in RAxML using a phylogenetic constraint (Manos, 2016; McVay *et al*., 2017b; Hipp *et al*., 2018) available in the supplemental methods and supplemental data. The remainder of the tree was unconstrained and conforms closely to previous topologies.

We utilized neighbor-net (Bryant & Moulton, 2004) to visualize overall patterns of molecular genetic diversity. Likelihood-based methods (e.g., Solís-Lemus & Ané, 2016; Solís-Lemus *et al*., 2017; Wen *et al*., 2018; Zhang *et al*., 2018) that we have utilized on smaller oak datasets (Crowl *et al*., In review; Eaton *et al*., 2015; Hauser *et al*., 2017; McVay *et al*., 2017b,a) proved computationally intractable for the current dataset. Consequently, we utilized a splits network inferred with SPLITSTREE v. 14.3 (Huson & Bryant, 2006) based on the maximum-likelihood (GTR+gamma) pairwise distance matrix estimated in RAxML and the same datasets utilized for the singletons tree. Full phylogenetic analysis details are in the Supplement (Methods S1).

### Calibration of singletons tree

Branch lengths on the tree were inferred using penalized likelihood under both a relaxed model, where rates are uncorrelated among branches (Paradis, 2013), and a correlated rates model (which corresponds to the penalized likelihood approach of Sanderson, 2002), as implemented in the chronos function of ape v 5.1 (Paradis *et al*., 2004) of R v 3.4.4 (“Someone to Lean On”) (R-Development-Core-Team, 2004). Nodes were calibrated in two different ways, either using eight fossil calibrations, corresponding to the crown of the genus and seven key clades (Fig. S2a; Table 1), or more conservatively as stem ages, using a subset of five fossils (Fig. S2b; Table 1). The two calibrations (referred to as the ‘crown calibration’ and ‘stem calibration’ respectively) bracket what we consider to be plausible age ranges for the tree. A separate estimate of the best fit λ for the correlated clock model was made using cross-validation as implemented in the chronopl function of ape, and that value of λ was used for both the relaxed and correlated clocks. Comparison of □IC was used to identify the best fit model for each value of λ. Analysis details are in the Supplement (Methods S1)

**Table 1.** Fossil calibrations used in this study, with nodes indicated as most recent common ancestor of selected taxa. Max and min indicate maximum and minimum ages for calibrations in Ma. Crown calibration node and stem calibration node indicate the taxa whose MRCA are the calibration points for the crown and stem calibration analyses respectively. References cited in Table S2.

| Node | Max | Min | Crown calibration node | Stem calibration node |
| --- | --- | --- | --- | --- |
| Quercus – genus | 56 | 56 | Quercus | Quercus\|Notholithocarpus |
| section Lobatae | 47.87 | 47.87 | Quercus_agrifolia\|Quercus_emoryi | Quercus_agrifolia\|Quercus_arizonica |
| section Cyclobalanopsis | 48.32 | 48.32 | Quercus_gilva\|Quercus_acuta | Quercus_gilva\|Quercus_rehderiana |
| section Quercus | 45 | 45 | Quercus_lobata\|Quercus_arizonica | Quercus_pontica\|Quercus_arizonica |
| section Ilex | 47.8 | 37.8 | Quercus_franchetii\|Quercus_rehderiana |  |
| section Ilex – in part | 35.5 | 33.4 | Quercus_rehderiana\|Quercus_semecarpifolia |  |
| section Cerris – in part | 34 | 30 | Quercus_chenii\|Quercus_acutissima | Quercus_franchetii\|Quercus_cerris |
| section Cerris – European clade | 23 | 20.5 | Quercus_crenata\|Quercus_cerris |  |

Transitions in lineage diversification rates were estimated using the speciation-extinction model implemented in Bayesian Analysis of Macroevolutionary Mixtures (BAMM) (Rabosky, 2014); the BAMMtools R package was used for configuration and analysis of MCMC. Priors were set using the setBAMMpriors function. Analyses were run for 4E06 generations, saving every 2000 generations, with four chains per MCMC analysis. To visualize changes in standing diversity over time for the different sections, we plotted lineage through time (LTT) plots by section against δ^18^O levels reported in Zachos *et al*. (2001) as a temperature proxy. Analysis details are in the Supplement (Methods S1).

### Investigating the genomic landscape of oak evolutionary history

Introgressive status of loci for two known introgression events involving the Eurasian white oaks (McVay *et al*., 2017b) and the western North American lobed-leaf white oaks (McVay *et al*., 2017a) was assessed by calculating the likelihood of phylogenies inferred for each locus under the constraint of the inferred divergence history (species tree) and the gene flow history at odds with that divergence history, as inferred in the studies cited above. These two cases are of particular interest because they are well studied, and lineage sorting has been ruled out in the above studies as an explanation of incongruence between the alternative topologies we test. Position of loci with a relative support of at least 2 log-likelihood points for one history relative to the other were mapped back to the *Quercus robur* genome (Plomion *et al*., 2018). Analysis details are in the Supplement (Methods S1).

To identify relative phylogenetic informativeness of loci, two tests were conducted based on the singletons tree. First, the ML topology was estimated in RAxML for each of 2,762 mapped, rootable loci of at least 10 individuals that resolved at least one bipartition. Overall, locus trees resolved an average of 4.48 (+/−1.83 s.d.) nodes, with a maximum of 15 and a median of 4. These were compared with the total-evidence tree using quartet similarities using the tqDist algorithm (Sand *et al*., 2014) in the Quartet package (Smith, 2019). We used as our similarity metric the number of quartets resolved the same way for both the locus tree and the whole singletons tree divided by the sum of quartets resolved the same or differently. Then, these same locus trees were mapped back to the singletons tree using phyparts (Smith *et al*., 2015), which identifies for all branches on a single tree how many individual locus trees support or reject that branch. We tested for genomic autocorrelation in phylogenetic signal using spline correlograms (Bjørnstad & Falck, 2001; Bjørnstad, 2008), with each chromosome tested independently. Analysis details are in the Supplement (Methods S1).

## Results

### RAD-seq data matrix

RAD-seq library preps and sequencing yielded a mean of 1.685E06 ± 1.104E06 (s.d.) raw reads per individual; of these, > 99.8% (1.683E06 ± 1.104E06) passed quality filters. The total number of clusters per individual prior to clustering across individuals was 101,895 ± 58,810, with a mean depth of 17.2 ± 11.2 sequences per individual and cluster. Clusters with more than 10,000 sequences per individual were discarded. Mean estimated heterozygosity by individual was 0.0135 ± 0.0027, and sequencing error rate was 0.0020 ± 0.0004. After clustering, a total of 49,991 loci were present in at least 15 individuals each. Each individual in the final dataset posseses 6.48% ± 2.48% of all clustered loci. The total data matrix is 4.352 × 10^6^ aligned nucleotides in width. The singletons dataset is composed of 22,432 loci present in at least 15 individuals, making up a dataset of 1.970 × 10^6^ aligned nucleotides.

### All-tips tree

The all-tips tree (Fig. S1) comprises 246 named *Quercus* species, of which 99 have a single sample. The remaining 147 species have an average of 3.54 ± 2.72 (s.d.) samples each. 97 of the 147 species with more than one sample cohere for all samples, and only 13 have a taxonomic disparity index (TDI, Pham *et al*., 2016) of 10 or more (Table S3), suggesting taxonomic problems beyond difficulties distinguishing very close relatives. All but four are Mexican species or species split between the southwestern U.S. and Mexico (see Discussion). Of the others, the largest TDI values are for *Q. stellata* and *Q. parvula* of North America, *Q. hartwissiana* and *Q. petraea* of western Eurasia, all with a complicated taxonomic history.

The topology of the all-tips tree closely matches previous analyses based on fewer taxa (McVay *et al*., 2017b; Hipp *et al*., 2018; Deng *et al*., 2018) for all sections except sections *Quercus* and *Virentes*. Unlike previous analyses, the all-tips topology embeds the long-branched section *Virentes* within section *Quercus*, sister to a clade comprising the SW US and Mexican clade and the Stellatae clade. This appears to be an artefact of clustering, as prior analyses of the same taxa do not reveal this topology, and unconstrained analysis of these taxa also recovers this aberrant topology. As a consequence, we consider the large-scale topology of the white oaks *s.l.* not to be reliable in the all-tips tree, and as this topology is well resolved in prior works (McVay *et al*., 2017b,a), we constrain the singletons topology as described in the methods section.

### Topology and timing of the oak phylogeny

Between the correlated and relaxed models of molecular rate heterogeneity, the correlated rates model (i.e., the penalized likelihood approach of Sanderson 2002) is consistently favored using □IC except at λ of 0, when the models are identical (Table S4). Though dating estimates differ little from λ = 0 to λ = 10 (not shown, but reproducible using code archived for this paper), cross-validation shows lowest sensitivity of taxon-removal on dating estimates at λ = 1.

Analyses with the crown-age calibrations (Fig. 1, S3a) suggest an older origin of most sections than proposed in prior studies (e.g., Cavender-Bares *et al*., 2015; Hipp *et al*., 2018; Deng *et al*., 2018), in part because in the current study we had access to a more comprehensive picture of the fossil record in oaks, including fossils used as age priors that predate those used in earlier studies. Section *Virentes* in our analysis has a crown age of ca. 30 Ma, whereas Cavender-Bares *et al*. (2015) estimated the crown age at 11 Ma. Even under the stem-age calibrations (Fig. 1; Fig. S3b, c), we estimate the crown age of *Virentes* at close to the Oligocene-Miocene boundary (ca. 23 Ma), nearly twice as old as prior estimates. Sections *Quercus* and *Lobatae* had an Oligocene crown constraint (31 Ma) in our previous work (Hipp *et al*., 2018); in the current study, they were constrained to a mid-Eocene origin (45–48 Ma) for the crown calibration, while the stem calibration recovers a late-Eocene origin for the red oaks (39 Ma) while the white oaks float down to a mid-Oligocene crown age (28 Ma). In the previous study of section *Cyclobalanopsis*, a minimum age of 33 Ma was set as a constraint at the root of subgenus *Cerris*, leading to a late Oligocene crown age for section *Cyclobalanopsis* (Deng *et al*., 2018); by contrast we recover an early Eocene crown age (38 Ma) for the group under the crown calibration, late Eocene (36 Ma) under the stem calibration. Given the high fossil density in *Quercus* (Table 1 and references therein; also reviewed in part in Denk & Grimm, 2009; Grímsson *et al*., 2015; Denk *et al*., 2017), the potential for alternative interpretations of their placement, and disparity among alternative methods for modeling (Paradis, 2013; Donoghue Philip C. J. & Yang Ziheng, 2016), we leave an investigation of a broader range of dating scenarios to later studies.

**Fig. 1.**
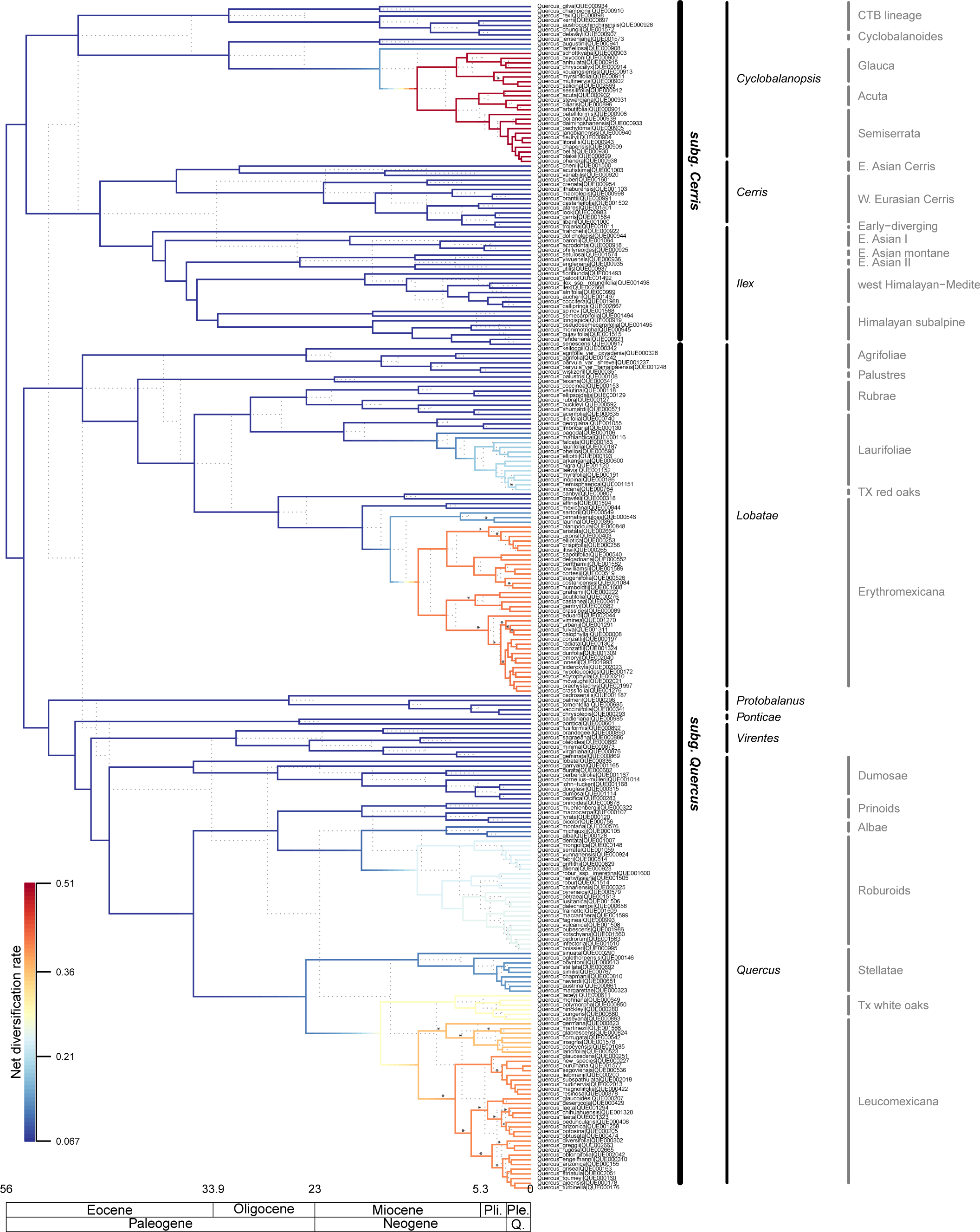
Singletons tree, calibrated using eight crown calibration fossils (solid lines) or 5 stem-calibration fossils (dotted lines). Single exemplars per species were analyzed using maximum likelihood; multiple samples are included for some species to represent cryptic or undescribed diversity (e.g., in *Quercus arizonica*, *Q. laeta*, *Q. conzattii*) or named infraspecies (e.g., varieties of *Quercus agrifolia* and *Q. parvula*). Labels to the right of the tree indicate subgenera (black) and sections (medium gray) following the latest taxonomy for the genus (Denk *et al*., 2017). Branch colors represent net diversification rates estimated using reversible-jump MCMC in BAMM (Rabosky, 2014), integrating over uncertainty in the timing and location of shifts in lineage diversification rates. rjMCMC was conducted with explicit lineage-specific sampling proportions specified, and thus accounts for the relatively low species sampling in the Mexican / Central American oaks and the southeast Asian section *Cyclobalanopsis*. All bootstrap values > 100 except for nodes marked with *, which are all 80-99 except for two: the common ancestor of *Q. costaricensis* and *Q. humboldtii* and the MRCA of *Q. myrsinifolia* and *Q. salicina* both have bootstrap values < 5.

White oaks *s.str.* are estimated in the crown-calibration analysis to have arrived in Eurasia some point in the Oligocene, close to the split between the section *Ponticae* sisters, which despite their morphological similarity appear to have diverged from one another nearly twice as long ago as the crown age of the Mexican white oaks; under the stem-calibration, the Eurasian white oaks are approximately half the crown age of the *Ponticae*. By contrast with the two species of sect. *Ponticae*, the Mexican white oak ancestor gave rise to an estimated 80 species in approximately half the time. The Roburoids had divided into a European and an East Asian clade by the early Miocene under the crown calibration, the late Miocene under the stem calibration.

Under the diversification scenarios implied by both the crown and the stem calibrations (Fig. 1, 2), there are four relatively recent and nearly simultaneous upticks in diversification: white oaks of Mexico and Central America; the red oaks of Mexico and Central America; the Eurasian (Roburoid) white oaks; and the Glauca, Semiserrata, and Acuta clades of section *Cyclobalanopsis*. In addition, the Eurasian white oaks and the southeastern U.S. white oaks (the Stellatae clade) and red oaks (the Laurofoliae clade) show a lesser increase in diversification rate in both analyses, and the clade of section *Ilex* that includes the Himalayan and Mediterranean species shows an uptick in diversification rate in the stem calibration. This result is robust to missing taxa, as we find essentially the same clades increasing in rate even assuming the 40% of missing taxa in our study were missing at random from the tree (Fig. S3a-c), with the addition of a portion of section *Ilex* and some of the eastern North American taxa as high-rate clades under the global sampling proportions model.

**Fig. 2.**
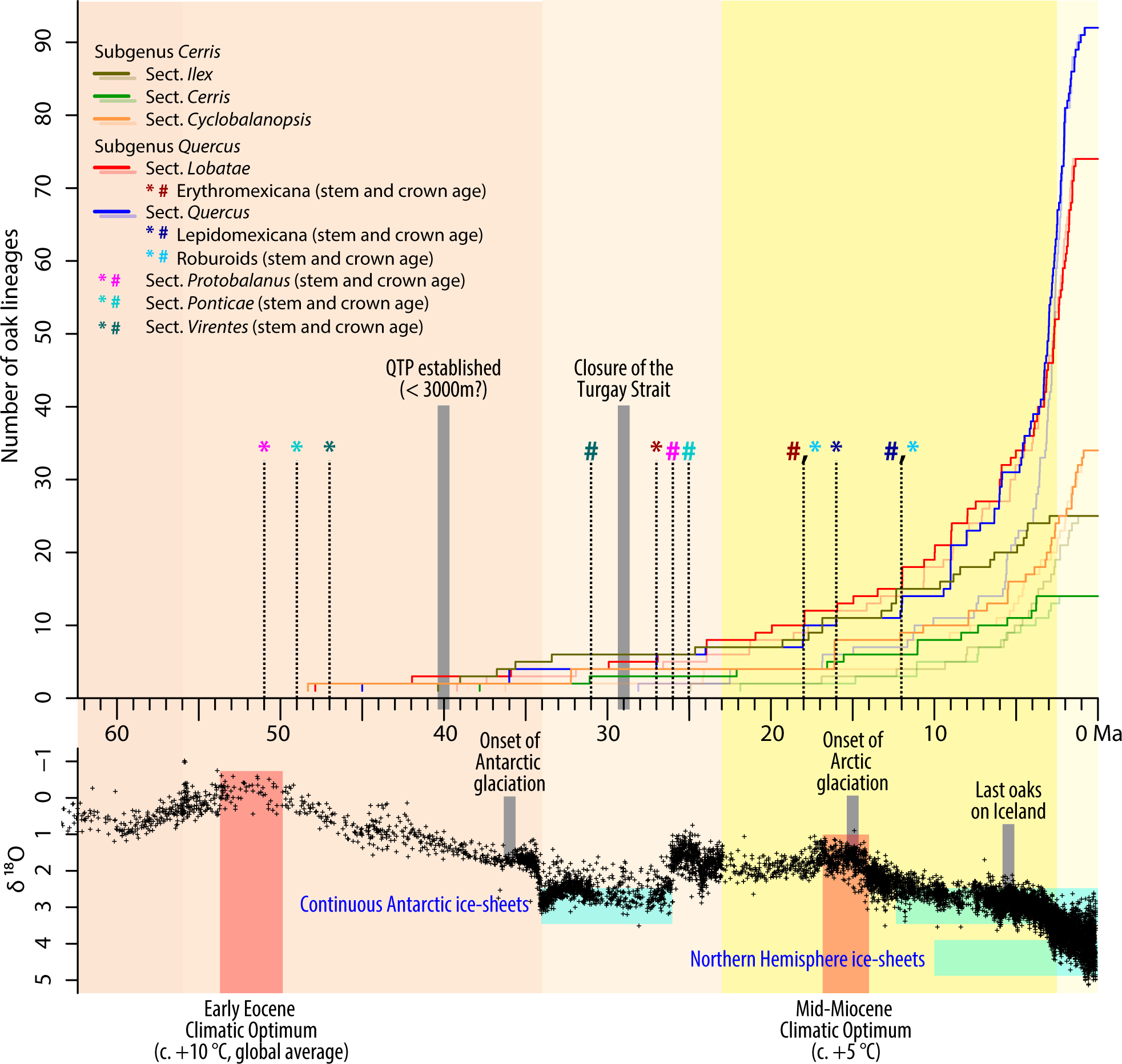
Lineages-through-time (LTT) plot showing the diversification of five species-rich lineages (sections *Cerris, Cyclobalanopsis, Ilex, Lobatae, Quercus*) within genus *Quercus* for the preferred (early fossils treated as crown group representatives) and conservative dating (dimmed lines, fossils treated as stem group taxa). Major tectonic events on the Northern Hemisphere (formation of the Qinghai-Tibetan Plateau, QTP (Scotese, 2014; Botsyun *et al.*, 2019); closure of the Turgai Sea) and global climate context (based on marine stable isotope data; Zachos *et al.*, 2001) shown for comparison. Timing of onset of Arctic glaciation and viability of the North Atlantic Land Bridge for oak migration are reviewed in Denk *et al*. (2010, 2013)and literature therein. Background colors indicate Cenozoic epochs/periods (following Walker *et al*. (2018); from left to right: Paleocene, Eocene, Oligocene, Miocene, Quaternary).

### Genomic arrangement of RAD-seq loci

A total of 39,860 loci aligned to at least one position on the oak genome. The 12 “pseudochromosomes” (inferred linkage groups, corresponding to the 12 *Quercus* chromosomes) as well as 360 scaffolds that did not map to the linkage groups were targeted by these loci. A total of 19,468 loci mapped to a unique position on a scaffold placed to one of the 12 oak genome pseudochromosomes, an average of 1,622.3 ± 575.4 (s.d.) per chromosomes. Of these, 31.7% ± 8.1% overlapped with the boundaries of a gene model (Fig. 3), despite the fact that only 10.1% of the 716 Mb of the *Quercus robur* genome that fall within the 12 pseudochromosomes fall within the endpoints of a gene model.

**Fig. 3.**
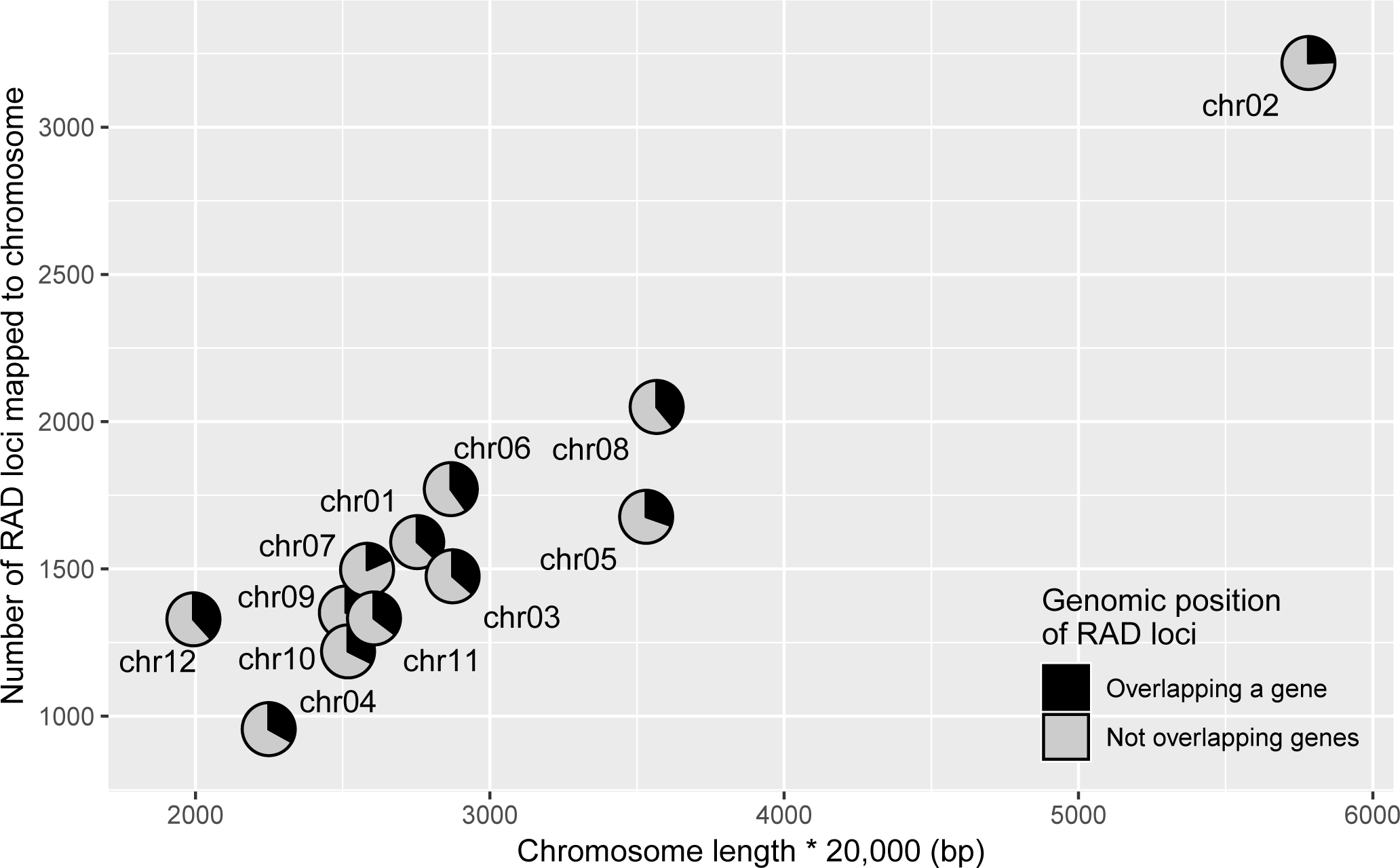
RAD-seq loci by chromosome. RAD-seq loci mapping to a unique position on one of the twelve *Quercus robur* genome pseudochromosomes are included in this figure and analyses reported in the paper. Chromosome length is based on total sequence length of scaffolds assigned to the *Q. robur* pseudochromosomes. Genomic position of loci overlapping vs not overlapping a gene was determined by detecting overlap of the RAD-seq locus start and end points with start and end points of the 25,808 gene models reported for the *Q. robur* genome (Plomion *et al.*, 2018).

For the tests of introgression, 2,422 loci had taxon sampling appropriate to testing for introgression involving *Q. macrocarpa* and *Q. lobata* (the Dumosae alternative topologies); 2,228 were suitable to testing for introgression involving the Roburoid white oaks and *Q. pontica* (the Roburoid alternative topologies); and 728 were suitable to testing both. Because we were interested in investigating genomic overlap in support for different areas of the species tree, we limited ourselves to the 728 loci that were potentially informative about both situations. Of these, 418 mapped to one position on one of the *Quercus robur* pseudochromosomes; and of these, 297 exhibited a log-likelihood difference of at least 2.0 between the better and more poorly supported topology for the Dumosae hypothesis or the Roburoid hypothesis, or both (Fig. 4). There was no correlation between the Roburoid and Dumosae hypotheses (*r* = −0.0286, *p* = 0.4878), meaning that loci that support or reject either of the Roburoid hypotheses do not correlate with a particular Dumosae hypothesis. Moreover, whether or not a locus is located within one of the *Q. robur* gene models has no effect on whether it recovers the introgression or the divergence history for the Roburoid oaks (*F*_1,366_ = 0.6494, *p* = 0.4209) or the Dumosae (*F*_1,415_ = 0.0377, *p* = 0.8461).

**Fig. 4.**
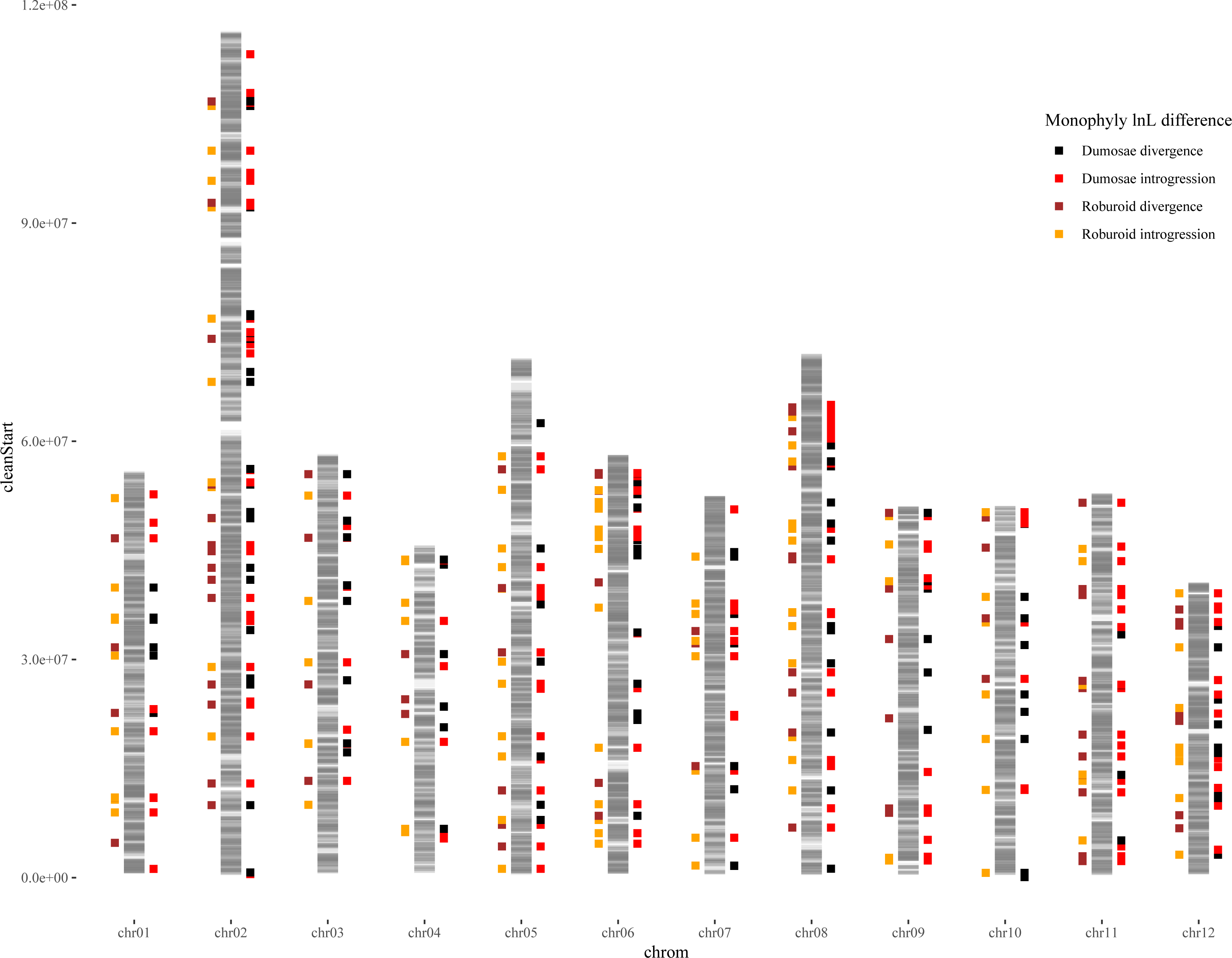
Genomic distribution of loci favoring alternative placements of the Roburoid white oaks and of *Quercus lobata* / *Quercus macrocarpa*. The 19,468 RAD-seq loci that map to a single position on one of the *Quercus robur* pseudochromosomes are represented by gray bands; chromosomal areas of darker gray have a denser mapping of RAD-seq loci. Mapped beside the chromosomes are the positions of 325 RAD-seq loci with a log-likelihood difference of at least 2 between trees constrained to be monophyletic for the Roburoids vs those placing the Roburoids with *Q. pontica* (194 loci); those differing by at least 2 between trees constrained to be monophyletic for both the Dumosae and the Prinoids vs those placing *Q. lobata* or *Q. macrocarpa* in the opposing clade (290 loci); or both (159 loci). These two hypotheses were selected because the topological differences have been demonstrated in prior studies (Crowl *et al.*, In review; McVay et al., 2017b,a) to be a consequence of introgression, not lineage sorting alone. The relative mapping of these loci thus provides a study in the distribution of loci that are informative about population divergence history vs. ancient introgression in two closely related clades. The mismatch between loci (*r* = −0.0286, *p* = 0.4878) suggests that introgression is not genomically conserved.

Quartet similarity—the number of taxon quartets with a topology shared between trees over the total number of quartets that both trees are informative about—between the RAD-seq individual-locus trees and the singletons tree (Fig. S4) is similarly uninfluenced by presence in one of the gene models presented in the *Quercus robur* genome (Plomion *et al*., 2018) (*F*_1,2542_ = 0.0495, *p* = 0.8239) and shows no evidence of genomic auto-correlation (Fig. S5). Rather, loci that support the tree are distributed across the genome. The same is true using locus trees to investigate the support for selected nodes of the phylogeny, all strongly supported (bootstrap support > 95% for all nodes tested; Fig. S1) (Fig. 5). The 2762 RAD-seq locus trees made 4,745 branch-level support claims and 27,283 conflict claims on the singletons tree, of which 6,409 total claims pertain to the nodes investigated, ranging from 107 to 1,055 per node (427.3 ± 273.7; Fig. 5). The locus-by-locus incongruence is high at this level: the proportion of loci concordant with each node averages 0.2395 ± 0.2523, but the range is high, from 0.6879 for the genus as a whole to as low as 0.0075 for the Mexican red oaks and 0.0088 for the Mexican white oaks (Table S5). There is no genomic autocorrelation in support vs. rejection of nodes in the singletons tree by individual locus trees (as inferred using phyparts; Smith *et al*., 2015) (Fig. S6), but the correlation between the crown age of clades investigated and the proportion of loci concordant with the crown age is positive and moderately significant (*r* = 0.4996, *p* = 0.0579; Fig. S7). Three clades stand out as outliers for high proportion of loci supporting divergence (outside the 95% regression CI): the genus as a whole, and sections *Cerris* and *Ilex*. This widespread genomic incongruence is reflected in broad network-like reticulation in the neighbor-net tree at the base of most clades (Fig. 6).

**Fig. 5.**
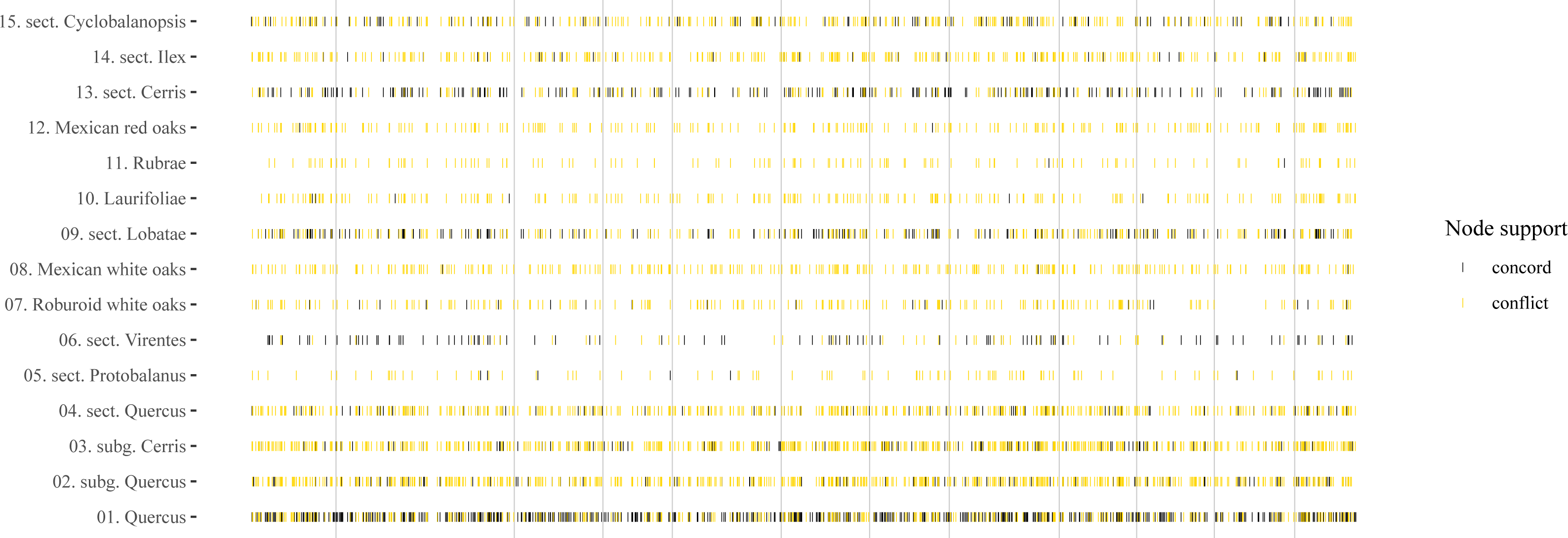
Loci congruent vs. discordant with key nodes of phylogeny. An average of 123.9 (± 178.9) RAD-seq locus trees are informative about each of the 15 named clades represented in this figure. Dark bands indicate RAD-seq loci that support a node; light bands indicate loci that conflict with it.

**Fig. 6.**
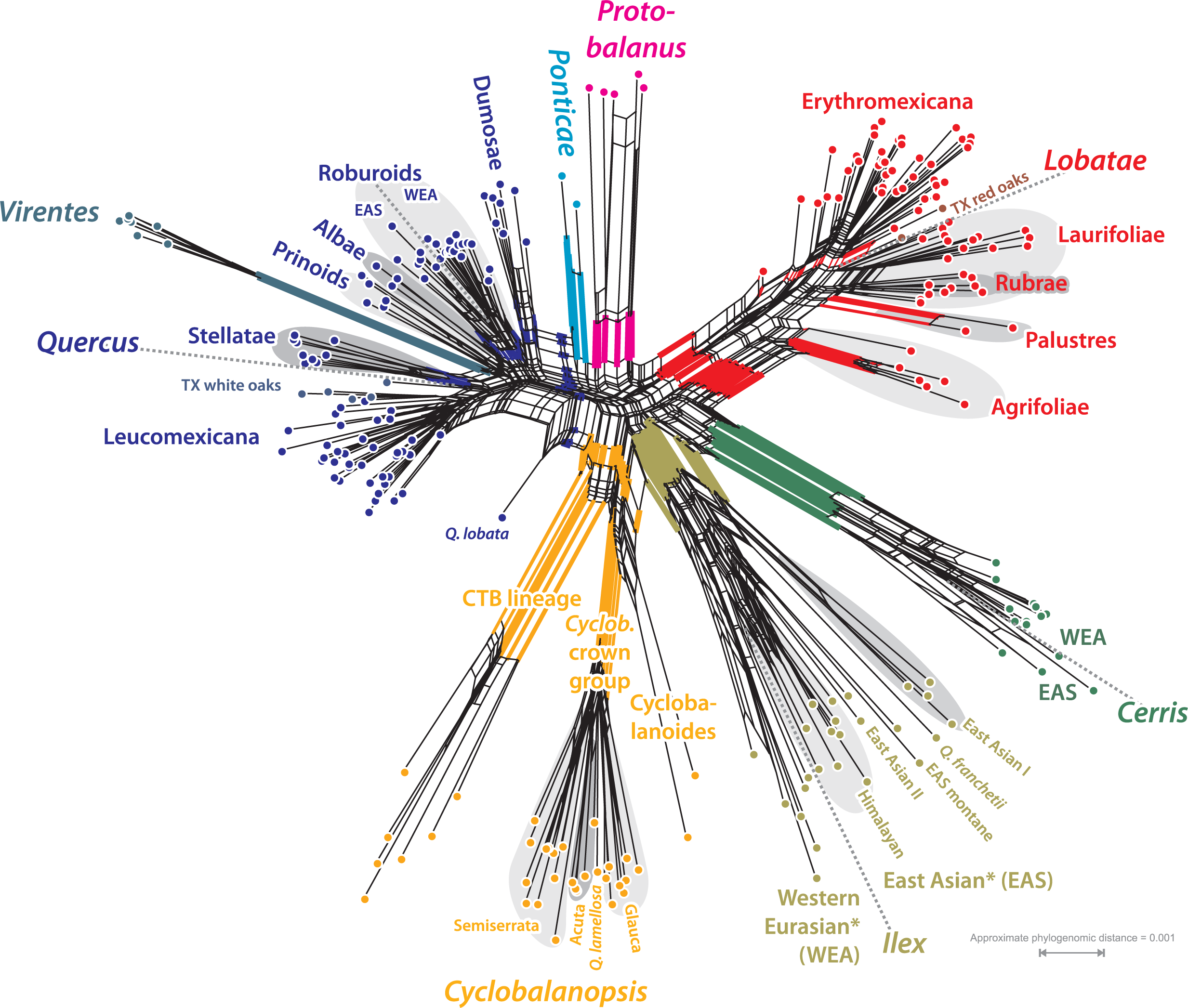
Neighbor-net, planar (meta-)phylogenetic network based on pairwise ML distances. Members of the major clades with unambiguous (tree) support (cf. Fig. 1) are clustered. All currently accepted sections are color-coded; edge bundles defining neighborhoods corresponding to sections and infra-sectional clades are colored accordingly. Main biogeographic splits within each section are indicated by dotted gray lines. The graph depicts the variance in inter- and intra-sectional genetic diversity patterns. The most genetically unique clades within each subgenus (sect. *Lobatae* for subgenus *Quercus*; sect. *Cerris* for subgenus *Cerris*) are placed on the right side of the graph; the distance to the spider-web-like center of the graph, which in this case may represent the point-of-origin (being also the mid-point between all tips and the connection of both subgenera) reflects the corresponding phylogenetic root-tip distances observed in the ML tree. Tree-like portions may be indicative of bottleneck situations in the formation of a clade; fan-like portions reflect potential genetic gradients developed during unhindered radiation (geographic expansion; note e.g. the position of Texan white and red oaks; strict West-East ordering within sect. *Ilex*), i.e. absence of major evolutionary bottlenecks.

## Discussion

Our analyses demonstrate that the diversity of oaks we observe today reflects deep geographic separation of major clades within the first 15 million years after the origin of the genus, and that standing species diversity arose mostly within the last 10 million years, predominantly in four rapidly diversifying clades that together account for ca. 60% of the diversity of the genus. Previous work has demonstrated American oak diversity was shaped in large part by ecological opportunity, first by the space left by tropical forests as they receded from North America, then by migration into the mountains of Mexico (Hipp *et al*., 2018; Cavender-Bares *et al*., 2018). The current study deepens this understanding by demonstrating two increases in diversification rates in Eurasia: one in the Eurasian white oaks, which arrived from eastern North America 7.5 to 18 Ma to low continental oak diversity, and no closely related oaks; and one in the southeast Asian section *Cyclobalanopsis*, driven by changing climates and the Himalayan orogeny (Deng *et al*., 2018). At the same time, our work demonstrates widespread genomic incongruence in phylogenetic history, with alternative phylogenetic histories interleaved across all linkage groups. Contrary to our hypothesis at the outset of this study, there appear to be no regions of the genome that on their own define the entire oak phylogeny. Instead, the primary divergence history of oaks (Crowl *et al*., In review; McVay *et al*., 2017b) knits together and emerges from a patchwork of histories that comprise the oak genome.

### Topology and timing of the global oak phylogeny

Our work indicates that by the mid-Eocene (45 Ma), all *Quercus* sections (*fide* Denk *et al*., 2017), representing eight major clades of the genus, had originated with the possible exception of section *Quercus*, which under the stem calibrations scenario arose at the Eocene-Oligocene boundary (33 Ma). Following this compressed interval of crown radiation, diversification rates spiked in the late Miocene to Pliocene, ca. 10 Ma (Fig. 2), primarily in southeast Asia, Mexico, and the white oaks of Eurasia. The eight fossil calibrations that we utilize here, and the two alternative methods of calibrating the tree (Fig. S3a-c), bracket what we consider to be a wide range of the plausible diversification times for the genus; so that while additional calibrations and a wider range of rate models bear investigation, we consider this overall finding for the shape and timing of oak diversification to be reasonable.

While *Quercus* arose at around the early Eocene climatic optimum (the earliest known *Quercus* fossil is pollen from Sankt Pankratz, Austria, 47°45’ N latitude, ca. 56 Ma; Hofmann *et al*., 2011), early fossils range as far north as Axel Heiberg Island in far northern Canada, which at 79° (both modern and paleolatitude in early Eocene; Scotese, 2014) is 20° further north than the northernmost oak populations today. As it followed the cooling climate southward, the genus remained largely a lineage of the northern temperate zone with some species of sections *Virentes*, *Lobatae*, and *Quercus* inhabiting tropical climates; but even these possess physiological adaptations that reflect their temperate ancestry (Cavender-Bares, 2019). In Eurasia, section *Cyclobalanopsis* dominates in subtropical evergreen broadleaf forests (Deng *et al*., 2018), but the sister sections *Cerris* and *Ilex* are temperate to Mediterranean. This climatic conservatism structures the geographic distribution of oak clades at several levels. Geographic patterns among and within major clades in the American oaks (subg. *Quercus*) have already been studied in detail, with geographic differentiation among the western U.S., the eastern U.S., and the southwestern U.S. and Mexico / Central America in each of two sections approximately simultaneously (Hipp *et al*., 2018). The current phylogeny makes clear that in the Eurasian white oaks of sect. *Quercus*, the Roburoid clade, the morphologically distinctive Mediterranean, dry-adapted species often treated as subsection *Galliferae* (e.g., Tschan & Denk, 2012) are distributed among all four subclades, suggesting that adaptations to the Mediterranean climate are convergent within the Roburoid clade; as discussed below under *Rapid diversification of the Eurasian white oaks*, it is geography rather than ecology or morphology that defines clades: species within clades are mostly separated by ecology, not geography. Likewise, the western Eurasian members of section *Ilex* form an inclusive subtree, in which the two widespread Mediterranean species *Q. coccifera* and *Q. ilex* are clearly separated and placed sister to the montane Asian clade. The geographically most distant species of the section are also genetically most distinct (Fig. 6). Even within clades, geographic structuring is evident. In section *Cerris*, for example, the east and west Eurasian species group in sister clades; within these latter, the western Mediterranean *Q. crenata* and *Q. suber* ‘corkish oaks’, the Near East ‘Aegilops’ oaks (*Q. brantii*, *Q. ithaburensis*, *Q. macrolepis*), and the remaining central-eastern Mediterranean members of the section are clearly separated. Within sect. *Quercus*, the North American Prinoids and Albae form a grade, reflecting diversification in North America predating dispersal of the Roburoid ancestor back to Eurasia. Once established in Eurasia, this lineage then diverged into East Asian and western Eurasian sister clades, ca. 10 My after isolation from its North American ancestors. Geography is imprinted in the oak phylogeny across clades, time periods, and continents.

Despite the older crown-age inferences in the current study in comparison to the RAD-seq studies of 2015–2018, relative dates in the present study confirm earlier results that the American oaks increased in diversification rate as they entered Mexico (in both red oaks and white oaks). It broadens this perspective with a global sample, providing evidence that the relative diversification rate of the Glauca, Acuta, and Semiserrata clades of the semitropical southeast Asian section *Cyclobalanopsis* is comparable to if not higher than the Mexican diversification. To a lesser extent, the Eurasian white oaks (the Roburoid clade) also show an increased rate of diversification. It is worth noting that the crown age of the Roburoid clade as a whole may be younger than our inferences, as fossil data raise some questions as to whether the Old World Roburoids were already isolated by the early Oligocene. Eocene sect. *Quercus* from Axel Heiberg Island (Canada), for example, appears to be closely allied with East Asian white oaks, and *Quercus furuhjelmi* from the Paleogene of Alaska and central Asia might belong to any of the modern New World or Old World white oak lineages, as might the early Oligocene *Quercus kodairae* and *Q. kobatakei* from Japan (Camus, 1936, 1938; Tanai & Uemura, 1994; Menitsky, 2005; Denk & Grimm, 2010; Tschan & Denk, 2012). Whereas previous analysis of *Fagus* (Fagaceae) found an unambiguous deep split between North American and Eurasian beech species that was also backed by fossils (Renner *et al*., 2016), the fossil data we have to date do not conclusively pin down the divergence between the North American and Eurasian white oaks. By contrast, the inferred early Miocene split between western Eurasian and East Asian white oaks is compatible with fossil evidence (Denk & Grimm, 2010), lending support to the observed increase in diversification rates observed in this study.

### Taxonomy of the Mexican and Central American oaks

The general high species-coherence we observe in the all-tips tree provides strong evidence that oak species, in general, are genetically coherent biological entities. The fact that 97 of the 147 species with more than one sample cohere for all samples provides the broadest test to date of species coherence in oaks. Among the species that do not exhibit coherence, the majority are from Mexico. Two sets of examples suggest that the Mexican oaks, while having been the focus of extensive taxonomic study (e.g., Trelease, 1924; Spellenberg & Bacon, 1996; Spellenberg *et al*., 1998; González-Villarreal, 2003; Valencia-A., 2004; de Beaulieu & Lamant, 2010), may harbor even higher species diversity than current estimates. The examples of *Quercus laeta* (González-Elizondo *et al*., In prep.) and *Q. conzattii* (McCauley *et al*., In revision; McCauley & Oyama, In prep.) exemplify a problem likely to be common in Mexican oaks. Both species have samples from northern and central to southern Mexico. Researchers working with them have noticed that northern and southern populations differ and may constitute separate species as our molecular data suggest. These samples are from two centers of Mexican oak diversity (Torres-Miranda *et al*., 2011, 2013; Rodríguez-Correa *et al*., 2015) and may reflect even higher species diversity in areas already known for high diversity. Interestingly, the observed divergence between northern Mexico and the Jalisco and Oaxaca samples in these examples appear to correlate with the formation of the Tepic-Zacoalco rift 5.5 Ma in the Jalisco block (Ferrari & Rosas-Elguera, 2000) and not with climatic transitions during the Pleistocene, which has been argued to be more a period of population movement than of speciation in the neotropics (Bennett *et al*., 2012). Notably, one of the youngest groups in the white oaks is located in the Sierra Madre Occidental, which harbors great habitat diversity in relatively small areas (Torres-Morales *et al*., 2010). The rugged and relatively young topography, a product of magmatism and subduction processes that lasted up through 12 Ma (Ferrari *et al*., 2018), and the convergence of temperate and tropical climates shaped the high diversification rates.

Several other cases of confusing taxonomy involving Mexican and Central American species are less clear. For example, the sect. *Lobatae* complex involving *Q. eugeniifolia*, *Q. benthamii*, *Q. cortesii* and *Q. lowilliamsii*, has a history of extensive taxonomic complication (Quezada Aguilar *et al*., 2016). The current work provides evidence that the species constitute a complex meriting more attention and draws attention to the possibility that Central American oak diversity and the role of Central American geology in Neotropical oak diversification has been underestimated (Cárdenes-Sandí *et al*., 2019), overshadowed as they have been by interest in the Mexican oak diversification (Quezada Aguilar *et al*., 2017). In the white oaks *s.str.* (sect. *Quercus*), cases such as *Q. insignis* and *Q. corrugata* seem even more obscure. Field observations (HG-C) suggest subtle differences between *Q. insignis*, a species of conservation concern from Jalisco, Oaxaca, Chiapas and Veracruz (Jerome, 2018), and *Q. corrugata* (from Chiapas and Oaxaca), but our molecular data are inconclusive. In general, taxonomy of the recently diverged or still-diverging Mexican species is particularly complicated because of extensive hybridization and introgression, even among relatively distantly related species (Spellenberg, 1995; Bacon & Spellenberg, 1996; González-Rodríguez *et al*., 2004; Bacon *et al*., 2011) and the dynamics of recent or ongoing speciation.

### Rapid diversification of Eurasian white oaks

Among the long-studied oaks of Eurasia (e.g., Camus, 1936, 1938, 1952; Schwarz, 1993; Menitsky, 2005), the data presented here point to the important role of ecological and morphological convergence among unrelated oaks. Phylogeny of the Eurasian white oaks (the Roburoid clade of section *Quercus*) has not previously been addressed in detail, despite their importance to our understanding of oak biodiversity and biology (cf. Kremer *et al*., 1991; Dumolin-Lapegue *et al*., 1997; Petit *et al*., 1997; Leroy *et al*., 2017 and references therein). Previous work has sampled a maximum of 14 Roburoid species (Hubert *et al*., 2014), but not recovered the monophyly of the clade, much less relationships among species. Our study includes 23 of the estimated 25 Roburoid white oak species, the strongest sampling to date. The late Miocene increase in diversification rate inferred in our study at the base of the western Eurasian white oaks clade is a particularly exciting finding, as it is one of only four major upticks in diversification inferred in our study. Our sampling of northern temperate white and red oaks is almost complete, and we have accounted for sampling bias in our diversification analyses, making it unlikely that the increase in diversification rate detected here is artefactual. The fact that the Roburoids are a northern temperate clade makes their radiation notable.

The unexpected increase in diversification rate in the Roburoids parallels the sympatric diversification of red and white oaks in North America, with divergence within clades and geographic regions accompanying convergence between clades (Cavender-Bares *et al*., 2018). As in the Mexican oak diversification (Torres-Miranda *et al*., 2011; Rodríguez-Correa *et al*., 2015), the western Eurasian white oaks are ecologically diverse, ranging from lowland swamp to Mediterranean scrub, steppe and from mesic lowland forests to subalpine timberline (de Beaulieu & Lamant, 2010). The European Roburoid clades are not readily diagnosable morphologically, and the morphological and ecological convergence among clades has led to taxonomic confusion. The morphologically distinctive Mediterranean, dry-adapted species (subsection *Galliferae*; cf. Tschan & Denk, 2012), for example, are distributed among all four subclades. Conversely, Roburoid clades 1 to 4 show geographic sorting whereas differentiation within clades commonly reflects ecological and climatic niche evolution along with morphological adaptations (e.g. from deciduous large lobed leaves to small, brevideciduous, unlobed leaves). Our study thus demonstrates that across the genus, ecological diversification within clades has shaped diversification.

### Genomic landscape of the global oak phylogeny

The current study uses mapped phylogenomic markers to demonstrate that the oak tree of life is etched broadly across the genome. Previous work demonstrated that approximately 19% of RAD-seq loci were associated with ESTs (Hipp *et al*., 2014), but that the EST-associated RAD-seq loci analyzed alone did not yield a topology that was different or differently supported than the RAD-seq loci not associated with EST markers, and that they were not differently apportioned to the base or the tips of the phylogeny (which might have suggested that RAD-seq loci associated with coding regions were more or less conservative or more or less homoplasious than the remainder). In the current study, 6,099 (31.3%) of RAD-seq loci in our dataset that map uniquely to one position in the genome do so in or overlapping with a predicted gene in the *Quercus robur* genome (as expected from a methylation-sensitive restriction enzyme; Rabinowicz *et al*., 2005; Pegadaraju *et al*., 2013). Our work demonstrates that gene-based RAD-seq loci do not differ from non-gene-based RAD-seq loci in similarity to the consensus tree or on introgression rates in the Roburoids and the Dumosae. Gene identity tells us little or nothing about how reliably a region of the genome records phylogenetic history.

At the same time, non-significant correlation between loci that strongly differentiate alternative topologies in the Dumosae and Roburoids suggests that these stories segregate nearly independently on the genome. There is also no evidence of genomic autocorrelation of phylogenetic informativeness in our study, despite the fact that our study has more mapped markers that significantly differentiate topologies in at least one of these parts of the tree than a previous study investigating genomic architecture of differentiation at the species level (*N* = 158 mapped markers with known G_ST_; Scotti-Saintagne *et al*., 2004). Our hypothesis that there are particular genes or regions of the genome that define the oak phylogeny globally appears to be false: rather, the phylogenetic history of oaks is defined by different genes in different lineages, making evolutionary history of oaks a phylogenetic and genomic mosaic. The effort to find a single best suite of genes for phylogenetic or population genetic inference across the oak genus is thus unlikely to be successful, though markers can clearly be designed for individual clades (Guichoux *et al*., 2011; Fitzek *et al*., 2018). What is perhaps most remarkable is that this heterogeneity of histories covarying independently along the oak genome yields, in aggregate, an evolutionary history of the complex genus that mirrors the morphological and ecological diversity of living and fossil oak species.

## Conclusion

Questions about the genomic architecture of population differentiation and speciation are generally asked at fine scales (Leroy *et al*., 2017, 2018), at the point at which population level processes directly shape genomic differentiation. But microevolution—comprising processes at the population level— leaves an imprint in the phylogeny; when such impressions persist, they can often be detected using topological methods that may be sensitive even to introgression along internal phylogenetic branches (Eaton *et al*., 2015; Solís-Lemus & Ané, 2016; McVay *et al*., 2017b). With multiple Fagaceae genomes now becoming available (Staton *et al*., 2015; Plomion *et al*., 2016, 2018; Sork *et al*., 2016; Ramos *et al*., 2018), we may soon be able to detangle the mosaic history of oaks and understand what story each gene tells. The current study makes clear that the phylogeny we unravel will neither be unitary nor told by a small subset of the genome, as the regions of the genome capturing the divergence history for one clade are not the regions capturing the divergence history of another. Understanding phylogenetic history in the face of this variation is only one problem. It will be followed by a greater one: how do we interpret the history of oak diversification in space and time if it is really a collection of diverse histories from different regions of the genome, all reflecting different evolutionary pathways, all equally real?

## Supporting information

Combined supplements -- all except large files

Table S1 -- Sample table

Fig S1 -- all-tips phylogeny, multiple pages

## Acknowledgements

Funding for this project was provided by U.S. National Science Foundation Awards 1146488 to ALH, 1146102 to PSM and 1146380 to JCB; Swedish Research Grants 2015-03986 to TD and 2009-00000 to GWG; support of the German Centre for Integrative Biodiversity Research (iDiv) Halle-Jena-Leipzig funded by the German Research Foundation (FZT 118) to E-DS and GWG; and The Morton Arboretum Center for Tree Science. This paper is dedicated to the memory of Michael Avishai (1935-2018), founder of the Jerusalem Botanical Gardens and cherished colleague.

## Author Contributions

ALH, PSM, JCB, MD, AK, CP, and AG-R conceived and designed the study.ALH, PSM, MH, MA, JCB, MD, TD, OG, MSG-E, AG-R, GWG, X-LJ, JDM, HR-C, MCS, VLS, and SV-A collected, identified, and curated samples. ALH, PSM, MH, JCB, AC, MD, TD, AG-R, GWG, X-LJ, JDM, VLS generated and analyzed phylogenetic data. CB, AK, IL, CP generated and analyzed genomic data. ALH, PSM, TD and GWG drafted the manuscript. All authors wrote and edited the manuscript.

## Supplement

**Fig. S1.** All-tips tree split by page (separate PDF)

**Fig. S2a.** Fossil calibration points: crown calibrations

**Fig. S2b.** Fossil calibration points: stem calibrations

**Fig. S3a.** Crown calibrations, global sampling estimate (60%)

**Fig. S3b.** Stem calibrations with rates, assuming clade-specific sampling proportions.

**Fig. S3c.** Stem calibrations, global sampling estimate (60%)

**Fig. S4.** Quartet similarity between individual loci and the full, all-tips tree, mapped to chromosomes

**Fig. S5.** Splines by chromosomes – quartets

**Fig. S6.** Splines by chromosomes – phyparts

**Fig. S7.** Phypart components

**Table S1.** Sampling table (separate XLSX)

**Table S2.** Citations for fossil calibrations.

**Table S3.** Taxonomic disparity index (TDI) for all unique species

**Table S4.** □IC values for alternative calibrations

**Table S5**. Phypart components and clade ages

**Methods S1.** Analysis details.

## References

Anderson E. 1953. Introgressive Hybridization. Biological Reviews 28: 280–307.

Bacon JR, Dávila-Aranda PD, Spellenberg R, González-Elizondo MS. 2011. The taxonomic status of the Mexican oak *Quercus undata* (Fagaceae, *Quercus*, section *Quercus*). Revista Mexicana de Biodiversidad 82: 1123–1131.

Bacon JR, Spellenberg R. 1996. Hybridization in two distantly related Mexican black oaks *Quercus conzattii* and *Quercus eduardii* (Fagaceae: *Quercus*: section *Lobatae*). SIDA, Contributions to Botany 17: 17–41.

Baird NA, Etter PD, Atwood TS, Currey MC, Shiver AL, Lewis ZA, Selker EU, Cresko WA, Johnson EA. 2008. Rapid SNP Discovery and Genetic Mapping Using Sequenced RAD Markers. PLOS ONE 3: e3376.

de Beaulieu A le H, Lamant T. 2010. Guide Illustré des Chênes. Geer, Belgium: Edilens.

Bennett K, Bhagwat S, Willis K. 2012. Neotropical refugia. The Holocene 22: 1207–1214.

Bjørnstad ON. 2008. ncf: spatial nonparametric covariance functions.

Bjørnstad ON, Falck W. 2001. Nonparametric spatial covariance functions: Estimation and testing. Environmental and Ecological Statistics 8: 53–70.

Boecklen WJ. 2017. Topology of syngameons. Ecology and Evolution 7: 10486–10491.

Botsyun S, Sepulchre P, Donnadieu Y, Risi C, Licht A, Rugenstein JKC. 2019. Revised paleoaltimetry data show low Tibetan Plateau elevation during the Eocene. Science 363: eaaq1436.

Bryant D, Moulton V. 2004. Neighbor-Net: An Agglomerative Method for the Construction of Phylogenetic Networks. Molecular Biology and Evolution 21: 255–265.

Camacho C, Coulouris G, Avagyan V, Ma N, Papadopoulos J, Bealer K, Madden TL. 2009. BLAST+: architecture and applications. BMC Bioinformatics 10: 421.

Camus AA. 1936. Monographie du genre Quercus. Tome I. Genre Quercus. Sous-genre Cyclobalanopsis et sous-genre Euquercus (Section Cerris et Mesobalanu). Paris: Editions Paul Lechevalier.

Camus AA. 1938. Monographie du genre Quercus. Tome II. Genre Quercus. Sous-genre Euquercus (sections Lepidobalanus et Macrobalanus). Paris: Editions Paul Lechevalier.

Camus AA. 1952. Monographie du genre Quercus. Tome III. Genre Quercus. Sous-genre Euquercus (sections Protobalanus et Erythrobalanus). Paris: Editions Paul Lechevalier.

Cannon CH, Lerdau M. 2015. Variable mating behaviors and the maintenance of tropical biodiversity. Frontiers in Genetics 6.

Cannon CH, Scher CL. 2017. Exploring the potential of gametic reconstruction of parental genotypes by F1 hybrids as a bridge for rapid introgression. Genome 60: 713–719.

Cárdenes-Sandí GM, Shadik CR, Correa-Metrio A, Gosling WD, Cheddadi R, Bush MB. 2019. Central American climate and microrefugia: A view from the last interglacial. Quaternary Science Reviews 205: 224–233.

Cavender-Bares J. 2019. Diversification, adaptation, and community assembly of the American oaks (*Quercus*), a model clade for integrating ecology and evolution. New Phytologist 221: 669–692.

Cavender-Bares J, Gonzalez-Rodriguez A, Eaton DAR, Hipp AL, Beulke A, Manos PS. 2015. Phylogeny and biogeography of the American live oaks (*Quercus* subsection *Virentes*): A genomic and population genetics approach. Molecular Ecology 24: 3668–3687.

Cavender-Bares J, Kothari S, Meireles JE, Kaproth MA, Manos PS, Hipp AL. 2018. The role of diversification in community assembly of the oaks (*Quercus* L.) across the continental U.S. American Journal of Botany 105: 565–586.

Crowl AA, McVay JD, Manos PS, Hipp AL, Lemmon A, Lemmon E. In review. Uncovering the genomic signature of ancient introgression between white oak lineages (*Quercus*) using anchored enrichment. New Phytologist.

Deng M, Jiang X-L, Hipp AL, Manos PS, Hahn M. 2018. Phylogeny and biogeography of East Asian evergreen oaks (*Quercus* section *Cyclobalanopsis*; Fagaceae): Insights into the Cenozoic history of evergreen broad-leaved forests in subtropical Asia. Molecular Phylogenetics and Evolution 119: 170–181.

Denk T, Grimm GW. 2009. Significance of Pollen Characteristics for Infrageneric Classification and Phylogeny in *Quercus* (Fagaceae). International Journal of Plant Sciences 170: 926–940.

Denk T, Grimm GW. 2010. The oaks of western Eurasia: Traditional classifications and evidence from two nuclear markers. Taxon 59: 351–366.

Denk T, Grimm GW, Grímsson F, Zetter R. 2013. Evidence from ‘Köppen signatures’ of fossil plant assemblages for effective heat transport of Gulf Stream to subarctic North Atlantic during Miocene cooling. Biogeosciences 10: 7927–7942.

Denk T, Grimm GW, Manos PS, Deng M, Hipp AL. 2017. An Updated Infrageneric Classification of the Oaks: Review of Previous Taxonomic Schemes and Synthesis of Evolutionary Patterns. In: Tree Physiology. Oaks Physiological Ecology. Exploring the Functional Diversity of Genus Quercus L. Springer, Cham, 13–38.

Denk T, Grimsson F, Zetter R. 2010. Episodic migration of oaks to Iceland: Evidence for a North Atlantic ‘land bridge’ in the latest Miocene. American Journal of Botany 97: 276–287.

Dodd RS, Afzal-Rafii Z. 2004. Selection and dispersal in a multispecies oak hybrid zone. Evolution 58: 261–269.

Donoghue Philip C. J., Yang Ziheng. 2016. The evolution of methods for establishing evolutionary timescales. Philosophical Transactions of the Royal Society B: Biological Sciences 371: 20160020.

Dumolin-Lapegue S, Demesure B., Fineschi S, Come V. L, Petit RJ. 1997. Phylogeographic structure of white oaks throughout the European continent. Genetics 146: 1475–1487.

Eaton DAR. 2014. PyRAD: assembly of de novo RADseq loci for phylogenetic analyses. Bioinformatics (Oxford, England) 30: 1844–1849.

Eaton DAR, Hipp AL, González-Rodríguez A, Cavender-Bares J. 2015. Historical introgression among the American live oaks and the comparative nature of tests for introgression. Evolution 69: 2587–2601.

Edelman NB, Frandsen P, Miyagi M, Clavijo BJ, Davey J, Dikow R, Accinelli GG, Belleghem SV, Patterson NJ, Neafsey DE, et al. 2018. Genomic architecture and introgression shape a butterfly radiation. bioRxiv: 466292.

Ferrari L, Orozco-Esquivel T, Bryan SE, López-Martínez M, Silva-Fragoso A. 2018. Cenozoic magmatism and extension in western Mexico: Linking the Sierra Madre Occidental silicic large igneous province and the Comondú Group with the Gulf of California rift. Earth-Science Reviews 183: 115–152.

Ferrari L, Rosas-Elguera J. 2000. Late Miocene to Quaternary extension at the northern boundary of the Jalisco Block, western Mexico: The Tepic-Zacoalco Rift revised. In: Special Paper 334: Cenozoic tectonics and volcanism of Mexico. Geological Society of America, 41–63.

Fitzek E, Delcamp A, Guichoux E, Hahn M, Lobdell M, Hipp AL. 2018. A nuclear DNA barcode for eastern North American oaks and application to a study of hybridization in an Arboretum setting. Ecology and Evolution 8: 5837–5851.

Fitz-Gibbon S, Hipp AL, Pham KK, Manos PS, Sork V. 2017. Phylogenomic inferences from reference-mapped and de novo assembled short-read sequence data using RADseq sequencing of California white oaks (*Quercus* subgenus *Quercus*). Genome 60: 743–755.

Folk RA, Soltis PS, Soltis DE, Guralnick R. 2018. New prospects in the detection and comparative analysis of hybridization in the tree of life. American Journal of Botany 105: 364–375.

González-Rodríguez A, Arias DM, Valencia-A. S, Oyama K. 2004. Morphological and RAPD analysis of hybridization between *Quercus affinis* and *Q. laurina* (Fagaceae), two Mexican red oaks. American Journal of Botany 91: 401–409.

González-Villarreal LM. 2003. Two new species of oak (Fagaceae, *Quercus* sect. *Lobatae*) from the Sierra Madre del Sur, Mexico. Brittonia 55: 49–60.

Grímsson F, Zetter R, Grimm GW, Pedersen GK, Pedersen AK, Denk T. 2015. Fagaceae pollen from the early Cenozoic of West Greenland: revisiting Engler’s and Chaney’s Arcto-Tertiary hypotheses. Plant Systematics and Evolution 301: 809–832.

Guichoux E, Lagache L, Wagner S, Léger P, Petit RJ. 2011. Two highly validated multiplexes (12-plex and 8-plex) for species delimitation and parentage analysis in oaks (*Quercus* spp.). Molecular Ecology Resources 11: 578–585.

Hardin JW. 1975. Hybridization and introgression in *Quercus alba*. Journal of the Arnold Arboretum 56: 336–363.

Hauser DA, Keuter A, McVay JD, HIpp AL, Manos PS. 2017. The evolution and diversification of the red oaks of the California Floristic Province (*Quercus* section *Lobatae*, series *Agrifoliae*). American Journal of Botany 104: 1581–1595.

Hipp AL. 2017. American oaks in an evolutionary context. In: Jerome D, Beckman E, Kenny L, Wenzell K, Kua C-S, Westwood M, eds. The Red List of US Oaks. Lisle: USDA Forest Service and The Morton Arboretum, 10–11.

Hipp AL. 2018. Pharaoh’s Dance: the oak genomic mosaic. PeerJ Preprints 6: e27405v1.

Hipp AL, Eaton DAR, Cavender-Bares J, Fitzek E, Nipper R, Manos PS. 2014. A framework phylogeny of the American oak clade based on sequenced RAD data. PLoS ONE 9: e93975.

Hipp AL, Manos PS, González-Rodríguez A, Hahn M, Kaproth M, McVay JD, Avalos SV, Cavender-Bares J. 2018. Sympatric parallel diversification of major oak clades in the Americas and the origins of Mexican species diversity. New Phytologist 217: 439–452.

Hofmann C-C, Mohamed O, Egger H. 2011. A new terrestrial palynoflora from the Palaeocene/Eocene boundary in the northwestern Tethyan realm (St. Pankraz, Austria). Review of Palaeobotany and Palynology 166: 295–310.

Hubert F, Grimm GW, Jousselin E, Berry V, Franc A, Kremer A. 2014. Multiple nuclear genes stabilize the phylogenetic backbone of the genus *Quercus*. Systematics and Biodiversity 12: 405–423.

Huson DH, Bryant D. 2006. Application of Phylogenetic Networks in Evolutionary Studies. Molecular Biology and Evolution 23: 254–267.

Jerome D. 2018. Quercus insignis. IUCN Red List of Threatened Species 2018: e.T194177A2302931.

Jiang X-L, Deng M, Hipp AL, Su T, Yan M-X, Zho Z-K. In review. East Asian origins of European holly oaks via the Tibet-Himalayas. Journal of Biogeography.

Kim BY, Wei X, Fitz□Gibbon S, Lohmueller KE, Ortego J, Gugger PF, Sork VL. 2018. RADseq data reveal ancient, but not pervasive, introgression between Californian tree and scrub oak species (*Quercus* sect. *Quercus*: Fagaceae). Molecular Ecology 27: 4556–4571.

Kremer A, Petit R, Zanetto A, Fougère V, Ducousso A, Wagner D, Chauvin C. 1991. Nuclear and organelle gene diversity in *Quercus robur* and *Q. petraea*. In: Müller-Starck G, Ziehe M, eds. Genetic Variation in European Forest Trees. Frankfurt-am-Main: Sauerländer’s Verlag, 141–166.

Leroy T, Rougemont Q, Dupouey J-L, Bodenes C, Lalanne C, Belser C, Labadie K, Provost GL, Aury J-M, Kremer A, et al. 2018. Massive postglacial gene flow between European white oaks uncovered genes underlying species barriers. bioRxiv: 246637.

Leroy T, Roux C, Villate L, Bodénès C, Romiguier J, Paiva JAP, Dossat C, Aury J-M, Plomion C, Kremer A. 2017. Extensive recent secondary contacts between four European white oak species. New Phytologist 214: 865–878.

Lewis ZA, Shiver AL, Stiffler N, Miller MR, Johnson EA, Selker EU. 2007. High-Density Detection of Restriction-Site-Associated DNA Markers for Rapid Mapping of Mutated Loci in Neurospora. Genetics 177: 1163–1171.

Manos PS. 2016. Systematics and biogeography of the American oaks. International Oaks 27: 23–36.

McCauley RA, Cortés-Palomec AC, Oyama K. In revision. Species diversification in a lineage of Mexican red oak (Quercus section Lobatae subsection Racemiflorae) – the interplay between distance, habitat, and hybridization. Tree Genetics & Genomes.

McCauley RA, Oyama K. In prep. A re-evaluation of taxonomy in Quercus section Lobatae subsection Racemiflorae, resurrection of the name Q. pennivenia and description of a new taxon Q. huicholensis.

McVay JD, Hauser D, Hipp AL, Manos PS. 2017a. Phylogenomics reveals a complex evolutionary history of lobed-leaf white oaks in western North America. Genome 60: 733–742.

McVay JD, Hipp AL, Manos PS. 2017b. A genetic legacy of introgression confounds phylogeny and biogeography in oaks. Proc. R. Soc. B 284: 20170300.

Menitsky YL. 2005. Oaks of Asia.□: Science Publishers. Boca Raton: CRC Press.

Miller M, Atwood T, Eames BF, Eberhart J, Yan Y-L, Postlethwait J, Johnson E. 2007a. RAD marker microarrays enable rapid mapping of zebrafish mutations. Genome Biology 8: R105.

Miller MR, Dunham JP, Amores A, Cresko WA, Johnson EA. 2007b. Rapid and cost-effective polymorphism identification and genotyping using restriction site associated DNA (RAD) markers. Genome Research 17: 240–248.

Minh BQ, Nguyen MAT, von Haeseler A. 2013. Ultrafast Approximation for Phylogenetic Bootstrap. Molecular Biology and Evolution 30: 1188–1195.

Ortego J, Gugger PF, Sork VL. 2018. Genomic data reveal cryptic lineage diversification and introgression in Californian golden cup oaks (section *Protobalanus*). New Phytologist 218: 804–818.

Pääbo S. 2003. The mosaic that is our genome. Nature 421: 409–412.

Paradis E. 2013. Molecular dating of phylogenies by likelihood methods: A comparison of models and a new information criterion. Molecular Phylogenetics and Evolution 67: 436–444.

Paradis E, Claude J, Strimmer K. 2004. APE: Analyses of Phylogenetics and Evolution in R language. Bioinformatics 20: 289–290.

Pegadaraju V, Nipper R, Hulke B, Qi L, Schultz Q. 2013. De novo sequencing of sunflower genome for SNP discovery using RAD (Restriction site Associated DNA) approach. BMC Genomics 14: 556.

Petit R, Bodenes C, Ducousso A, Roussel G, Kremer A. 2003. Hybridization as a mechanism of invasion in oaks. New Phytologist 161: 151–164.

Petit RJ, Hampe A. 2006. Some Evolutionary Consequences of Being a Tree. Annual Review of Ecology, Evolution, and Systematics 37: 187–214.

Petit R, Pineau E, Demesure B, Bacilieri R, Ducousso A, Kremer A. 1997. Chloroplast DNA footprints of postglacial recolonization by oaks. Proceedings of the National Academy of Sciences USA 94: 9996–10001.

Pham KK, Hahn M, Lueders K, Brown BH, Bruederle LP, Bruhl JJ, Chung K-S, Derieg NJ, Escudero M, Ford BA, et al. 2016. Specimens at the Center: An Informatics Workflow and Toolkit for Specimen-Level Analysis of Public DNA Database Data. Systematic Botany 41: 529–539.

Pham KK, Hipp AL, Manos PS, Cronn RC. 2017. A Time and a Place for Everything: Phylogenetic history and geography as joint predictors of oak plastome phylogeny. Genome 60: 720–732.

Plomion C, Aury J-M, Amselem J, Alaeitabar T, Barbe V, Belser C, Bergès H, Bodénès C, Boudet N, Boury C, et al. 2016. Decoding the oak genome: public release of sequence data, assembly, annotation and publication strategies. Molecular Ecology Resources 16: 254–265.

Plomion C, Aury J-M, Amselem J, Leroy T, Murat F, Duplessis S, Faye S, Francillonne N, Labadie K, Provost GL, et al. 2018. Oak genome reveals facets of long lifespan. Nature Plants 4: 440–452.

Quammen D. 2018. The Tangled Tree: A Radical New History of Life. New York: Simon and Schuster.

Quezada Aguilar ML, Rodas-Duarte L, Marroquín-Tintí AA. 2016. Diversidad de encinos en Guatemala; una alternativa para bosques enegéticos, seguridad alimentaria y mitigación al cambio climático. Fase II. Jutiapa, Jalapa y Santa Rosa. Instituto de Investigaciones Químicas y Biológicas (IIQB). Centro de Estudios Conservacionistas (CECON), Facultad de Ciencias Químicas y Farmacia. Universidad de San Carlos de Guatemala.

Quezada Aguilar ML, Rodas-Duarte R, Marroquí-n-Tintí- AA. 2017. Contribución al conocimiento de los encinos (*Quercus*: Fagaceae) en los departamentos de Alta Verapaz, Baja Verapaz y Petén, Guatemala. Ciencia, Tecnologí-a y Salud 3: 115–126.

Rabinowicz PD, Citek R, Budiman MA, Nunberg A, Bedell JA, Lakey N, O’Shaughnessy AL, Nascimento LU, McCombie WR, Martienssen RA. 2005. Differential methylation of genes and repeats in land plants. Genome Research 15: 1431–1440.

Rabosky DL. 2014. Automatic Detection of Key Innovations, Rate Shifts, and Diversity-Dependence on Phylogenetic Trees. PLoS ONE 9: e89543.

Ramos AM, Usié A, Barbosa P, Barros PM, Capote T, Chaves I, Simões F, Abreu I, Carrasquinho I, Faro C, et al. 2018. The draft genome sequence of cork oak. Scientific Data 5: 180069.

R-Development-Core-Team. 2004. R: A language and environment for statistical computing. Vienna.

Ree RH, Hipp AL. 2015. Inferring phylogenetic history from restriction site associated DNA (RADseq). In: Hoerandl E, Appelhaus M, eds. Next Generation Sequencing in Plant Systematics. Koenigstein: Koeltz Scientific Books, 181–204.

Renner SS, Grimm GW, Kapli P, Denk T. 2016. Species relationships and divergence times in beeches: new insights from the inclusion of 53 young and old fossils in a birth–death clock model. Philosophical Transactions of the Royal Society B: Biological Sciences 371: 20150135.

Rodríguez-Correa H, Oyama K, MacGregor-Fors I, González-Rodríguez A. 2015. How Are Oaks Distributed in the Neotropics? A Perspective from Species Turnover, Areas of Endemism, and Climatic Niches. International Journal of Plant Sciences 176: 222–231.

Sand A, Holt MK, Johansen J, Brodal GS, Mailund T, Pedersen CNS. 2014. tqDist: a library for computing the quartet and triplet distances between binary or general trees. Bioinformatics 30: 2079–2080.

Sanderson MJ. 2002. Estimating absolute rates of molecular evolution and divergence times: a penalized likelihood approach. Molecular Biology and Evolution 19: 101–109.

Schwarz O. 1993. Quercus L. Flora Europaea I: 72–76.

Scotese CR. 2014. Atlas of Paleogene Paleogeographic Maps (Mollweide Projection), Maps 8-15, Volume 1, The Cenozoic, PALEOMAP Atlas for ArcGIS. Evanston, IL: PALEOMAP Project.

Scotti-Saintagne C, Mariette S, Porth I, Goicoechea PG, Barreneche T, Bodenes C, Burg K, Kremer A. 2004. Genome Scanning for Interspecific Differentiation Between Two Closely Related Oak Species [*Quercus robur* L. and *Q. petraea* (Matt.) Liebl.]. Genetics 168: 1615–1626.

Smith MR. 2019. Quartet: comparison of phylogenetic trees using quartet and bipartition measures. doi: 10.5281/zenodo.2536318.

Smith SA, Moore MJ, Brown JW, Yang Y. 2015. Analysis of phylogenomic datasets reveals conflict, concordance, and gene duplications with examples from animals and plants. BMC Evolutionary Biology 15: 150.

Solís-Lemus C, Ané C. 2016. Inferring Phylogenetic Networks with Maximum Pseudolikelihood under Incomplete Lineage Sorting. PLOS Genetics 12: e1005896.

Solís-Lemus C, Bastide P, Ané C. 2017. PhyloNetworks: A Package for Phylogenetic Networks. Molecular Biology and Evolution 34: 3292–3298.

Sork VL, Fitz-Gibbon ST, Puiu D, Crepeau M, Gugger PF, Sherman R, Stevens K, Langley CH, Pellegrini M, Salzberg SL. 2016. First Draft Assembly and Annotation of the Genome of a California Endemic Oak *Quercus lobata* Née (Fagaceae). G3: Genes, Genomes, Genetics 6: 3485–3495.

Spellenberg R. 1995. On the hybrid nature of *Quercus basaseachicensis* (Fagaceae, sect. *Quercus*). *Sida*, Contributions To Botany 16: 427–437.

Spellenberg R, Bacon JR. 1996. Taxonomy and Distribution of a Natural Group of Black Oaks of Mexico (*Quercus*, Section *Lobatae*, Subsection *Racemiflorae*). Systematic Botany 21: 85–99.

Spellenberg R, Bacon JR, González Elizondo MS. 1998. Los encinos (*Quercus*, Fagaceae) en un transecto sobre la Sierra Madre Occidentali. Boletín del Instituto de Botánica de la Universidad de Guadalajara 5: 357–387.

Stamatakis A. 2006. Phylogenetic models of rate heterogeneity: a high performance computing perspective. In: Proceedings 20th IEEE International Parallel Distributed Processing Symposium. 8 pp.

Stamatakis A. 2014. RAxML Version 8: A tool for Phylogenetic Analysis and Post-Analysis of Large Phylogenies. Bioinformatics 30: 1312–1313.

Staton M, Zhebentyayeva T, Olukolu B, Fang GC, Nelson D, Carlson JE, Abbott AG. 2015. Substantial genome synteny preservation among woody angiosperm species: comparative genomics of Chinese chestnut (*Castanea mollissima*) and plant reference genomes. BMC Genomics 16: 744.

Tanai T, Uemura K. 1994. Lobed oak leaves from the Tertiary of East Asia with reference to the oak phytogeography of the northern hemisphere. Transactions and proceedings of the Paleontological Society of Japan. New series 1994: 343–365.

Torres-Miranda A, Luna-Vega I, Oyama K. 2011. Conservation biogeography of red oaks (*Quercus*, section *Lobatae*) in Mexico and Central America. American Journal of Botany 98: 290–305.

Torres-Miranda A, Luna-Vega I, Oyama K. 2013. New Approaches to the Biogeography and Areas of Endemism of Red Oaks (*Quercus* L., Section *Lobatae*). Systematic Biology 62: 555–573.

Torres-Morales L, García-Mendoza DF, López-González C, Muñiz-Martínez R. 2010. Bats of Northwestern Durango, Mexico: Species Richness at the Interface of Two Biogeographic Regions. The Southwestern Naturalist 55: 347–362.

Trelease W. 1924. The American Oaks. Memoirs of the National Academy of Sciences 20: 1–255.

Tschan GF, Denk T. 2012. Trichome types, foliar indumentum and epicuticular wax in the Mediterranean gall oaks, *Quercus* subsection *Galliferae* (Fagaceae): implications for taxonomy, ecology and evolution. Botanical Journal of the Linnean Society 169: 611–644.

Valencia-A. S. 2004. Diversidad del género *Quercus* (Fagaceae) en México. Boletín de la Sociedad Botánica de México 75: 33–53.

Van Valen L. 1976. Ecological species, multispecies, and oaks. Taxon 25: 233–239.

Walker JD, Geissman JW, Bowring SA, Babcock LE. 2018. Geologic Time Scale v. 5.0: Geological Society of America, https://www.geosociety.org/GSA/Education_Careers/Geologic_Time_Scale/GSA/timescale/home.aspx.

Wen D, Yu Y, Zhu J, Nakhleh L. 2018. Inferring Phylogenetic Networks Using PhyloNet. Systematic Biology 67: 735–740.

Whittemore AT, Schaal BA. 1991. Interspecific gene flow in sympatric oaks. Proceedings of the National Academy of Sciences USA 88: 2540–2544.

Zachos J, Pagani M, Sloan L, Thomas E, Billups K. 2001. Trends, Rhythms, and Aberrations in Global Climate 65 Ma to Present. Science 292: 686–693.

Zhang C, Ogilvie HA, Drummond AJ, Stadler T. 2018. Bayesian Inference of Species Networks from Multilocus Sequence Data. Molecular Biology and Evolution 35: 504–517.

