## Supplementary material for "Genomic landscape of the global oak phylogeny": Combined supplements -- all except large files

### New Phytologist Supporting Information

Article title: **The genomic landscape of the global oak phylogeny**

Article acceptance date: Click here to enter a date.

The following Supporting Information is available for this article:

**Fig. S1** All-tips tree split by page (separate PDF)

**Fig. S2a** Fossil calibration points: crown calibrations

**Fig. S2b** Fossil calibration points: stem calibrations

**Fig. S3a** Crown calibrations, global sampling estimate (60%)

**Fig. S3b** Stem calibrations with rates, assuming clade-specific sampling propo

**Fig. S3c** Stem calibrations, global sampling estimate (60%)

**Fig. S4** Quartet similarity between individual loci and the full, all-tips tree, mapped to chromosomes

**Fig. S5** Splines by chromosomes - quartets

**Fig. S6** Splines by chromosomes - phypart

**Fig. S7** Phypart components

**Table S1** Sampling table (separate XLSX)

**Table S2** Citations for fossil calibrations

**Table S3** Taxonomic disparity index (TDI) for all unique species

**Table S4** PhiIC values for alternative calibrations

**Table S5** Phypart components and clade ages

**Methods S1** Analysis details

**Fig. S1** All-tips tree split by page (separate PDF)

**Fig. S2a** Fossil calibration points: crown calibrations

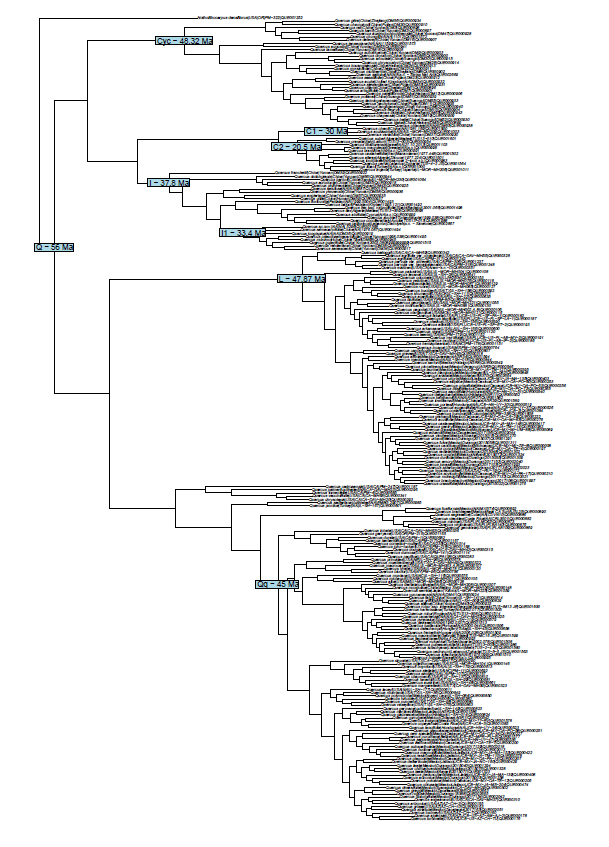

**Fig. S2b** Fossil calibration points: stem calibrations

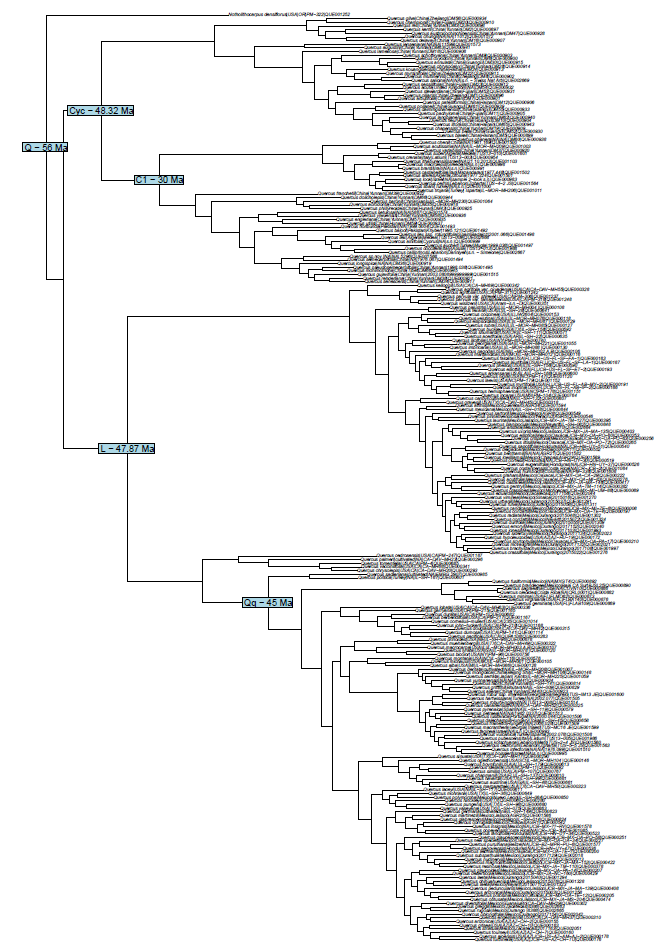

**Fig. S3a** Crown calibrations, global sampling estimate (60%)

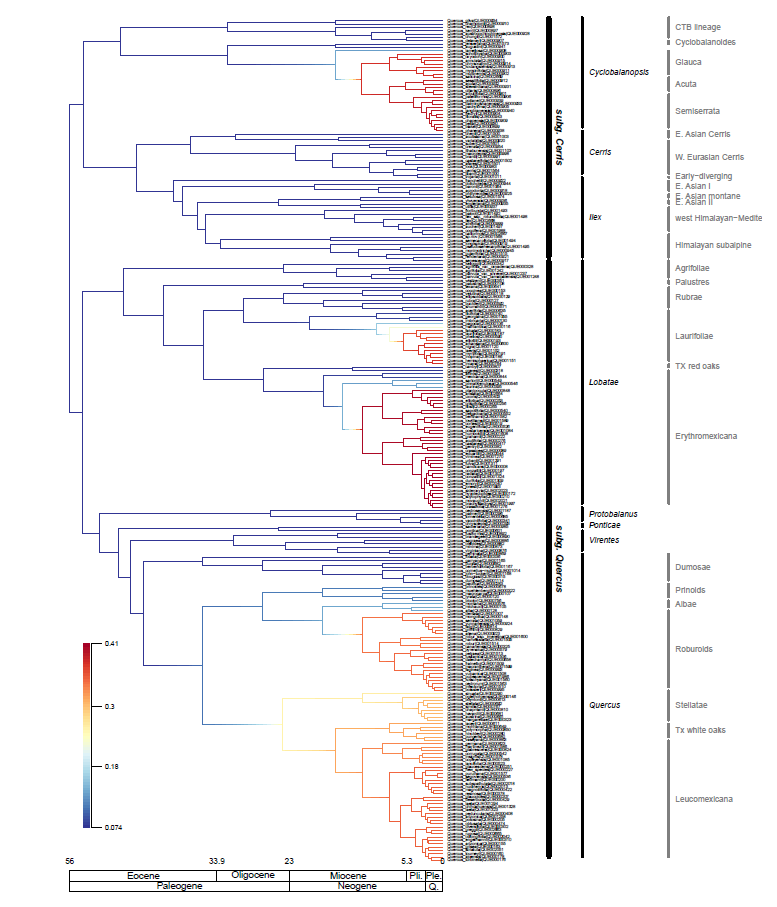

**Fig. S3b** Stem calibrations with rates, assuming clade-specific sampling proportions

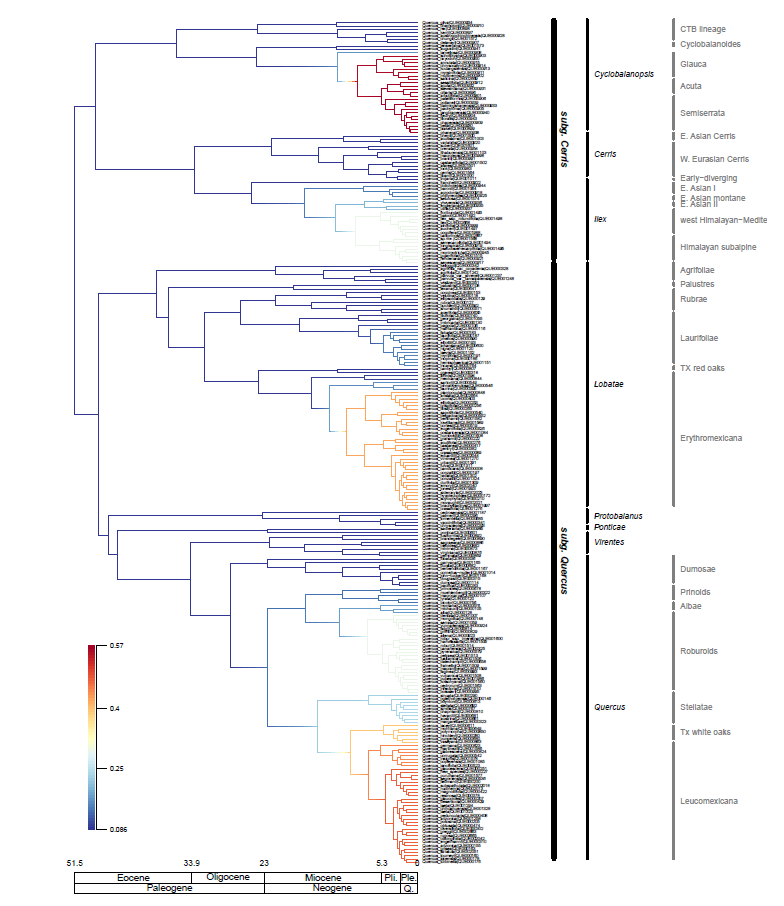

**Fig. S3c** Stem calibrations, global sampling estimate (60%)

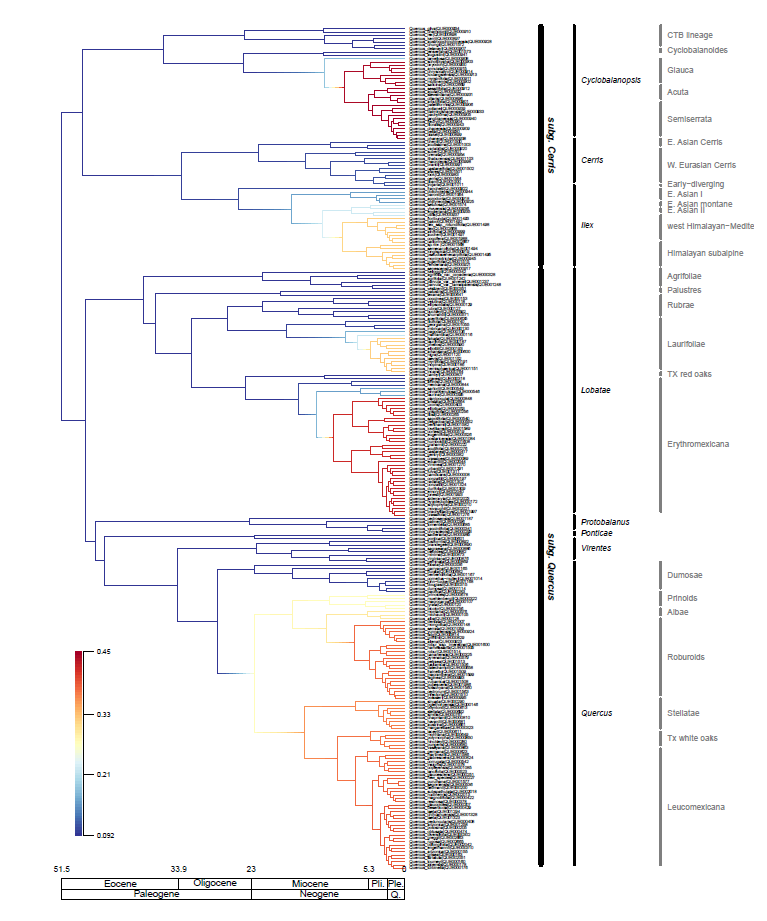

**Fig. S4** Quartet similarity between individual loci and the full, all-tips tree, mapped to chromosomes

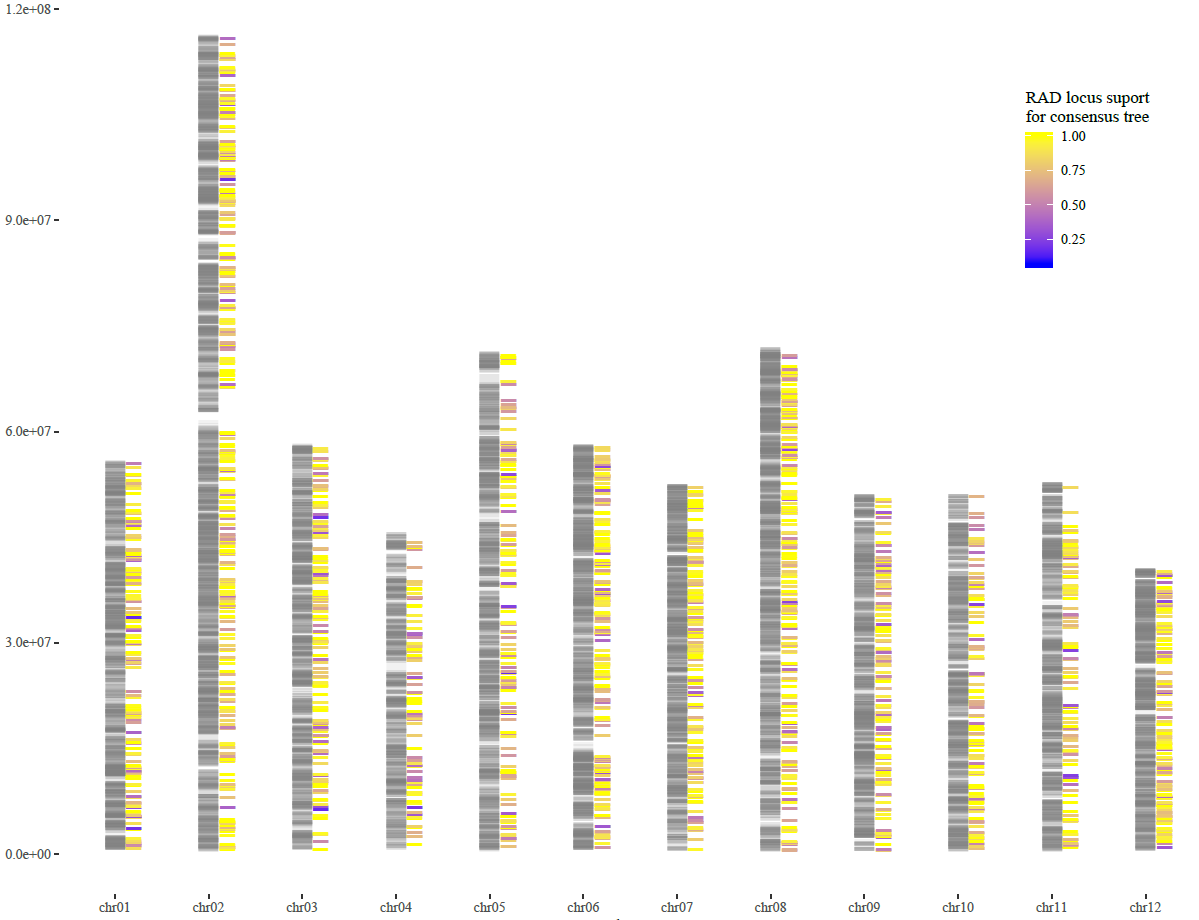

**Fig. S5** Splines by chromosomes – quartets. Spline correlograms indicate correlation in quartet scores for loci, plotted against the distance separating loci on the chromosome

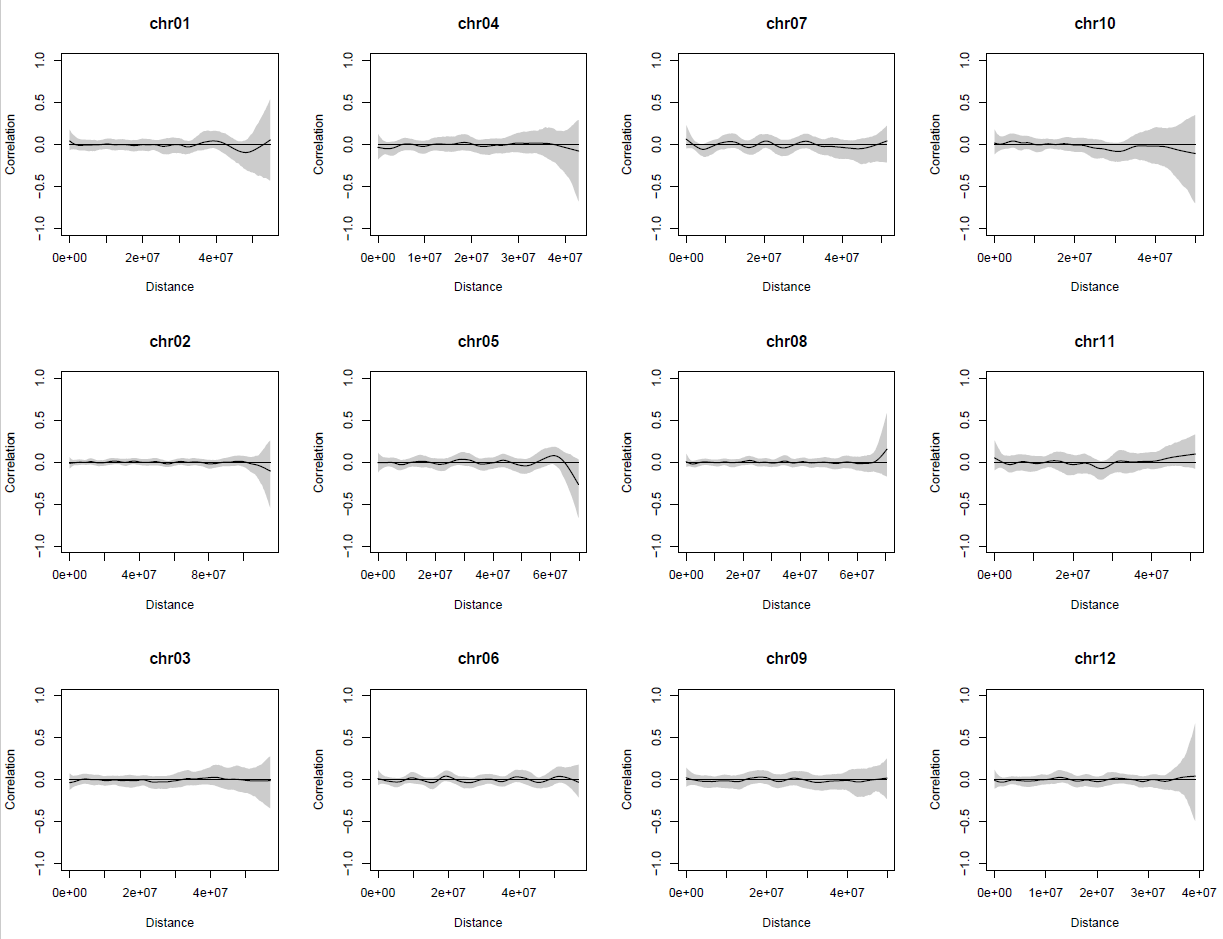

**Fig. S6** Splines by chromosomes – Phypart support (1) vs discord (0). Spline correlograms indicate correlation in Phypart support scores as binary values for all mapped loci, plotted against the distance separating loci on the chromosome.

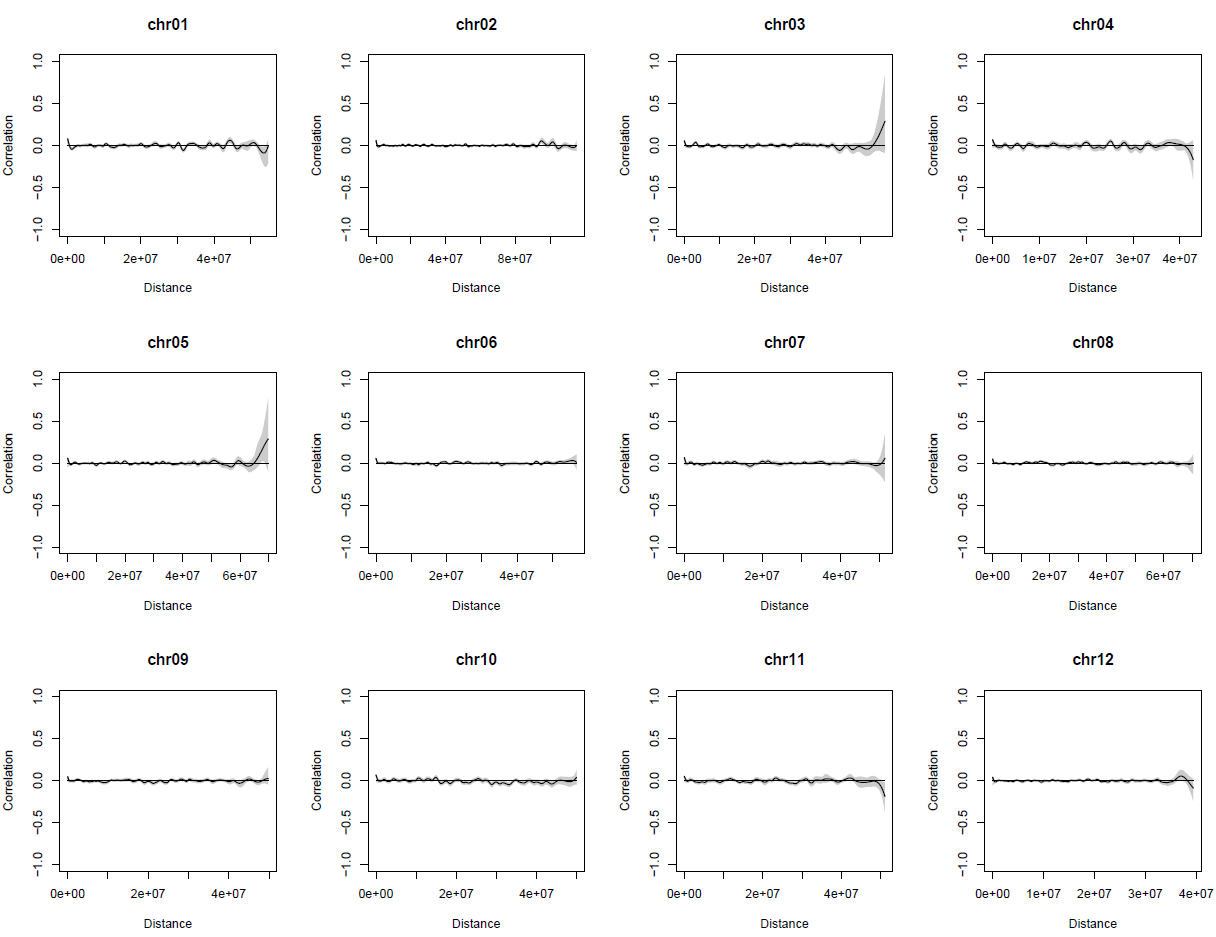

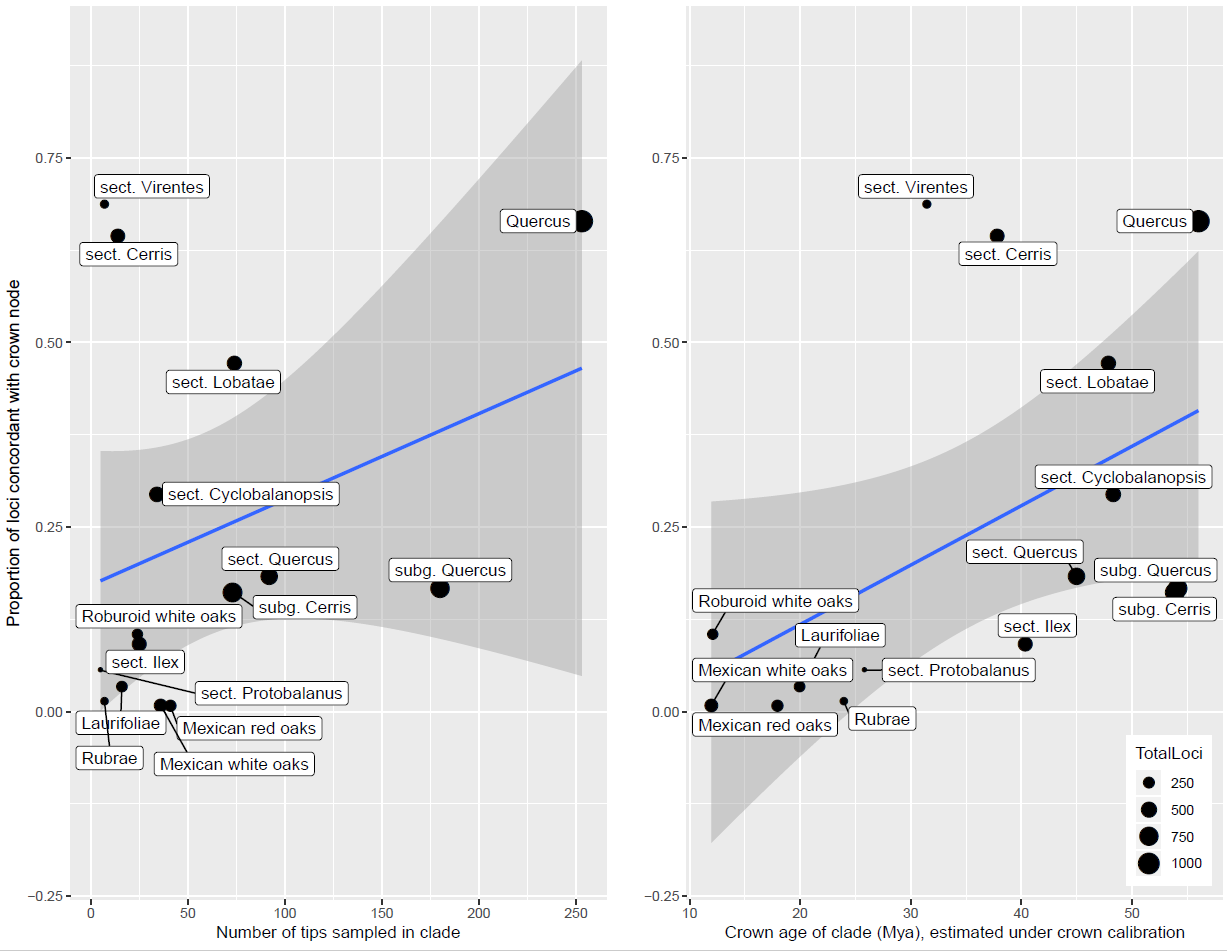
**Fig. S7** Phyparts components. Correlation between the crown age of clades investigated and the proportion of loci concordant with the crown age is positive and moderately significant (*r* = 0.4996, *p* = 0.0579). Three clades stand out as outliers for high proportion of loci supporting divergence (outside the 95% regression CI): the genus as a whole, and sections *Cerris* and *Ilex*. While age is significantly correlated with number of descendant tips sampled (*r* = 0.6623, *p* = 0.0052), there is no correlation between the proportion of loci concordant with a node and the number of sample in our dataset descendant from that node (*r* = 0.3251, *p* = 0.2371).

**Table S1** Sampling table (separate CSV file)

**Table S2** Fossil calibration notes and citations

| **node** | **citations** |
| --- | --- |
| *Quercus* – genus | *Quercus* pollen from the Paleocene/Eocene boundary (dated with dinoflagellates) |
| section *Lobatae* | Grímsson *et al.*, 2016: pollen of sect. *Lobatae* from the Princeton Chert (radiometric age 47.8 Ma, Greenwood et al., 2016) (Greenwood *et al.*, 2016; Grímsson *et al.*, 2016) |
| section *Cyclobalanopsis* | Manchester, 1994, 2011: fruits of sect. *Cyclobalanopsis* from the Clarno Formation, Nut Beds (radiometric age 48.32±0.11 Ma; Manchester 2011) (Manchester, 1994, 2011) |
| section *Quercus* | Interpreted to exclude sections *Ponticae* and *Virentes*. McIntyre, D.J., 1991; McIver & Basinger, 1999: Leaves from the forest swamp facies from the Geodetic Hills on Axel Heiberg Island stored at University of Saskatchewan Paleobotanical Collection (USPC) belong to sect. *Quercus*. Age is middle Eocene, c. 45 Ma (brontothere remains, Eberle & Greenwood, 2012). (McIntyre, D.J., 1991; McIver & Basinger, 1999; Eberle & Greenwood, 2012) |
| section *Ilex* | Hofmann, 2010: Pollen from the middle Eocene Changchang Formation, might be the oldest sect. *Ilex* pollen. Age (according to Spicer *et al.*, 2014), middle to late Eocene, i.e. 47.8-37.8 Ma. (Hofmann, 2010) |
| section *Ilex* – in part | Su et al., 2018: “Morphotype 22”, “Morphotype 25”, radiometric ages 34.6±0.8 – 35.5±0.3 Ma and 33.4±0.5 – 34.7±0.5 Ma. Leaf fossils belong to clade *Quercus semecarpifolia* to *Q. rehderiana*. (Su *et al.*, 2018) |
| section *Cerris* – in part | Pavlyutkin et al., 2015: *Quercus gracilis*, Kraskino Flora, Russian Far East, 34-30 Ma (correlation with adjacent radiometrically dated sites; Tanai & Uemura, 1994; Akhmetiev et al., 2009). Clade *Quercus chenii* to *Q. acutissima*. (Tanai & Uemura, 1994; Akhmetiev *et al.*, 2009; Pavlyutkin, 2015) |
| section *Cerris* – western Eurasian clade | Kmenta, 2011: Pollen, Altmittweida, Germany. 23-20.5 Ma (Standke et al., 2010). Western Eurasian clade of sect. *Cerris*. (Standke *et al.*, 2010; Kmenta, 2011) |

**Table S3** Taxonomic disparity index (TDI) for all unique species. Taxonomic disparity is calculated following Pham *et al.* (2016) as the difference between the number of tips in the clade defined by the most recent common ancestor (MRCA) of all individuals labeled as a given species and the number of tips actually labeled as that species. For *Quercus aliena*, for example, there are two individuals sequenced, but their MRCA defines a clade of 7 tips; thus their TDI = 5. Higher TDI indicates taxa that are more egregiously polyphyletic, species with a TDI ≥ 10 (see main text) are highlighted. The statistic is used heuristically to define taxa that are potentially problematic in the tree.

|  | Tips observed | MRCA clade | TDI (disparity) |
| --- | --- | --- | --- |
| *Quercus acerifolia* | 2 | 2 | 0 |
| *Quercus acrodonta* | 1 | 1 | 0 |
| *Quercus acuta* | 1 | 1 | 0 |
| *Quercus acutifolia* | 2 | 2 | 0 |
| *Quercus acutissima* | 1 | 1 | 0 |
| *Quercus afares* | 1 | 1 | 0 |
| *Quercus affinis* | 2 | 2 | 0 |
| *Quercus agrifolia* | 16 | 16 | 0 |
| *Quercus ajoensis* | 2 | 2 | 0 |
| *Quercus alba* | 5 | 5 | 0 |
| *Quercus aliena* | 2 | 7 | 5 |
| *Quercus alnifolia* | 1 | 1 | 0 |
| *Quercus annulata* | 1 | 1 | 0 |
| *Quercus arbutifolia* | 1 | 1 | 0 |
| *Quercus aristata* | 1 | 1 | 0 |
| *Quercus arizonica* | 11 | 24 | 13 |
| *Quercus arkansana* | 2 | 2 | 0 |
| *Quercus aucheri* | 1 | 1 | 0 |
| *Quercus augustini* | 1 | 1 | 0 |
| *Quercus austrina* | 2 | 2 | 0 |
| *Quercus austrocochinchinensis* | 1 | 1 | 0 |
| *Quercus baloot* | 1 | 1 | 0 |
| *Quercus baronii* | 1 | 1 | 0 |
| *Quercus bella* | 1 | 1 | 0 |
| *Quercus benthamii* | 1 | 1 | 0 |
| *Quercus berberidifolia* | 3 | 4 | 1 |
| *Quercus bicolor* | 4 | 4 | 0 |
| *Quercus blakei* | 1 | 1 | 0 |
| *Quercus boissieri* | 1 | 1 | 0 |
| *Quercus boyntonii* | 2 | 2 | 0 |
| *Quercus brachystachys* | 1 | 1 | 0 |
| *Quercus brandegeei* | 2 | 2 | 0 |
| *Quercus brantii* | 1 | 1 | 0 |
| *Quercus buckleyi* | 2 | 2 | 0 |
| *Quercus calliprinos* | 4 | 4 | 0 |
| *Quercus calophylla* | 2 | 2 | 0 |
| *Quercus canariensis* | 1 | 1 | 0 |
| *Quercus canbyi* | 2 | 2 | 0 |
| *Quercus castanea* | 4 | 15 | 11 |
| *Quercus castaneifolia* | 1 | 1 | 0 |
| *Quercus cedrorum* | 2 | 5 | 3 |
| *Quercus cedrosensis* | 1 | 1 | 0 |
| *Quercus cerris* | 6 | 8 | 2 |
| *Quercus championii* | 1 | 1 | 0 |
| *Quercus chapensis* | 1 | 1 | 0 |
| *Quercus chapmanii* | 2 | 2 | 0 |
| *Quercus chenii* | 1 | 1 | 0 |
| *Quercus chihuahuensis* | 3 | 3 | 0 |
| *Quercus chrysocalyx* | 1 | 1 | 0 |
| *Quercus chrysolepis* | 3 | 8 | 5 |
| *Quercus chungii* | 1 | 1 | 0 |
| *Quercus ciliaris* | 1 | 1 | 0 |
| *Quercus coccifera* | 3 | 7 | 4 |
| *Quercus coccinea* | 3 | 3 | 0 |
| *Quercus conzattii* | 6 | 15 | 9 |
| *Quercus copeyensis* | 1 | 1 | 0 |
| *Quercus cornelius-mulleri* | 3 | 3 | 0 |
| *Quercus corrugata* | 2 | 4 | 2 |
| *Quercus cortesii* | 3 | 6 | 3 |
| *Quercus costaricensis* | 2 | 2 | 0 |
| *Quercus crassifolia* | 3 | 5 | 2 |
| *Quercus crassipes* | 2 | 2 | 0 |
| *Quercus crenata* | 2 | 5 | 3 |
| *Quercus crispifolia* | 1 | 1 | 0 |
| *Quercus daimingshanensis* | 1 | 1 | 0 |
| *Quercus dalechampii* | 1 | 1 | 0 |
| *Quercus delavayi* | 1 | 1 | 0 |
| *Quercus delgadoana* | 1 | 1 | 0 |
| *Quercus dentata* | 1 | 1 | 0 |
| *Quercus depressipes* | 1 | 1 | 0 |
| *Quercus deserticola* | 2 | 2 | 0 |
| *Quercus diversifolia* | 1 | 1 | 0 |
| *Quercus dolicholepis* | 2 | 2 | 0 |
| *Quercus douglasii* | 3 | 3 | 0 |
| *Quercus dumosa* | 1 | 1 | 0 |
| *Quercus durata* | 2 | 5 | 3 |
| *Quercus durifolia* | 2 | 2 | 0 |
| *Quercus eduardi* | 5 | 5 | 0 |
| *Quercus elliottii* | 2 | 2 | 0 |
| *Quercus ellipsoidalis* | 13 | 13 | 0 |
| *Quercus elliptica* | 5 | 5 | 0 |
| *Quercus emoryi* | 6 | 7 | 1 |
| *Quercus engelmannii* | 2 | 2 | 0 |
| *Quercus engleriana* | 1 | 1 | 0 |
| *Quercus eugeniifolia* | 2 | 8 | 6 |
| *Quercus fabri* | 1 | 1 | 0 |
| *Quercus faginea* | 2 | 2 | 0 |
| *Quercus falcata* | 2 | 2 | 0 |
| *Quercus fleuryi* | 1 | 1 | 0 |
| *Quercus floribunda* | 1 | 1 | 0 |
| *Quercus frainetto* | 1 | 1 | 0 |
| *Quercus franchetii* | 1 | 1 | 0 |
| *Quercus fulva* | 2 | 2 | 0 |
| *Quercus fusiformis* | 4 | 6 | 2 |
| *Quercus gambelii* | 8 | 8 | 0 |
| *Quercus garryana* | 6 | 6 | 0 |
| *Quercus geminata* | 4 | 4 | 0 |
| *Quercus gentryi* | 5 | 5 | 0 |
| *Quercus georgiana* | 2 | 2 | 0 |
| *Quercus germana* | 4 | 4 | 0 |
| *Quercus gilva* | 1 | 1 | 0 |
| *Quercus glabrescens* | 1 | 1 | 0 |
| *Quercus glaucescens* | 1 | 1 | 0 |
| *Quercus glaucoides* | 3 | 3 | 0 |
| *Quercus grahamii* | 3 | 97 | 94 |
| *Quercus gravesii* | 1 | 1 | 0 |
| *Quercus greggii* | 3 | 14 | 11 |
| *Quercus griffithii* | 1 | 1 | 0 |
| *Quercus grisea* | 4 | 4 | 0 |
| *Quercus gujavifolia* | 1 | 1 | 0 |
| *Quercus hartwissiana* | 2 | 30 | 28 |
| *Quercus havardii* | 1 | 1 | 0 |
| *Quercus hemisphaerica* | 2 | 8 | 6 |
| *Quercus hinckleyi* | 1 | 1 | 0 |
| *Quercus humboldtii* | 1 | 1 | 0 |
| *Quercus hypoleucoides* | 2 | 2 | 0 |
| *Quercus ilex* | 3 | 3 | 0 |
| *Quercus ilicifolia* | 1 | 1 | 0 |
| *Quercus iltisii* | 1 | 1 | 0 |
| *Quercus imbricaria* | 2 | 2 | 0 |
| *Quercus incana* | 2 | 2 | 0 |
| *Quercus infectoria* | 2 | 5 | 3 |
| *Quercus inopina* | 2 | 2 | 0 |
| *Quercus insignis* | 2 | 4 | 2 |
| *Quercus ithaburensis* | 3 | 3 | 0 |
| *Quercus jenseniana* | 1 | 1 | 0 |
| *Quercus john-tuckeri* | 3 | 3 | 0 |
| *Quercus jonesii* | 3 | 3 | 0 |
| *Quercus kelloggii* | 13 | 13 | 0 |
| *Quercus kerrii* | 1 | 1 | 0 |
| *Quercus kotschyana* | 1 | 1 | 0 |
| *Quercus kouangsiensis* | 1 | 1 | 0 |
| *Quercus laceyi* | 2 | 2 | 0 |
| *Quercus laeta* | 7 | 48 | 41 |
| *Quercus laevis* | 2 | 2 | 0 |
| *Quercus lamellosa* | 1 | 1 | 0 |
| *Quercus lancifolia* | 2 | 2 | 0 |
| *Quercus langbianensis* | 1 | 1 | 0 |
| *Quercus laurifolia* | 2 | 2 | 0 |
| *Quercus laurina* | 2 | 4 | 2 |
| *Quercus libani* | 2 | 4 | 2 |
| *Quercus liebmanii* | 2 | 2 | 0 |
| *Quercus litoralis* | 1 | 1 | 0 |
| *Quercus lobata* | 3 | 3 | 0 |
| *Quercus longispica* | 1 | 1 | 0 |
| *Quercus look* | 1 | 1 | 0 |
| *Quercus lowilliamsii* | 2 | 4 | 2 |
| *Quercus lusitanica* | 1 | 1 | 0 |
| *Quercus lyrata* | 5 | 5 | 0 |
| *Quercus macranthera* | 2 | 2 | 0 |
| *Quercus macrocarpa* | 9 | 9 | 0 |
| *Quercus macrolepis* | 1 | 1 | 0 |
| *Quercus magnoliifolia* | 2 | 29 | 27 |
| *Quercus margarettae* | 2 | 4 | 2 |
| *Quercus marilandica* | 4 | 4 | 0 |
| *Quercus martinezii* | 2 | 2 | 0 |
| *Quercus mcvaughii* | 1 | 1 | 0 |
| *Quercus mexicana* | 2 | 2 | 0 |
| *Quercus michauxii* | 5 | 5 | 0 |
| *Quercus minima* | 4 | 8 | 4 |
| *Quercus mohriana* | 2 | 2 | 0 |
| *Quercus mongolica* | 1 | 1 | 0 |
| *Quercus monimotricha* | 1 | 1 | 0 |
| *Quercus montana* | 4 | 4 | 0 |
| *Quercus muehlenbergii* | 6 | 10 | 4 |
| *Quercus multinervis* | 1 | 1 | 0 |
| *Quercus myrsinifolia* | 1 | 1 | 0 |
| *Quercus myrtifolia* | 2 | 5 | 3 |
| *Quercus nigra* | 2 | 2 | 0 |
| *Quercus nudinervis* | 1 | 1 | 0 |
| *Quercus oblongifolia* | 4 | 4 | 0 |
| *Quercus obtusata* | 3 | 12 | 9 |
| *Quercus oglethorpensis* | 3 | 3 | 0 |
| *Quercus oleoides* | 5 | 6 | 1 |
| *Quercus oxyodon* | 1 | 1 | 0 |
| *Quercus pachyloma* | 1 | 1 | 0 |
| *Quercus pacifica* | 1 | 1 | 0 |
| *Quercus pagoda* | 2 | 2 | 0 |
| *Quercus palmeri* | 3 | 3 | 0 |
| *Quercus palustris* | 2 | 2 | 0 |
| *Quercus parvula* | 9 | 22 | 13 |
| *Quercus patelliformis* | 1 | 1 | 0 |
| *Quercus peduncularis* | 2 | 2 | 0 |
| *Quercus petraea* | 4 | 22 | 18 |
| *Quercus phanera* | 1 | 1 | 0 |
| *Quercus phellos* | 2 | 2 | 0 |
| *Quercus phillyreoides* | 1 | 1 | 0 |
| *Quercus pinnativenulosa* | 2 | 2 | 0 |
| *Quercus planipocula* | 1 | 1 | 0 |
| *Quercus poilanei* | 1 | 1 | 0 |
| *Quercus polymorpha* | 2 | 2 | 0 |
| *Quercus pontica* | 4 | 4 | 0 |
| *Quercus potosina* | 5 | 77 | 72 |
| *Quercus prinoides* | 4 | 8 | 4 |
| *Quercus pseudosemecarpifolia* | 1 | 1 | 0 |
| *Quercus pubescens* | 2 | 2 | 0 |
| *Quercus pungens* | 2 | 2 | 0 |
| *Quercus purulhana* | 2 | 2 | 0 |
| *Quercus pyrenaica* | 1 | 1 | 0 |
| *Quercus radiata* | 3 | 3 | 0 |
| *Quercus rehderiana* | 1 | 1 | 0 |
| *Quercus resinosa* | 4 | 29 | 25 |
| *Quercus rex* | 1 | 1 | 0 |
| *Quercus robur* | 6 | 7 | 1 |
| *Quercus rubra* | 19 | 19 | 0 |
| *Quercus rugosa* | 7 | 14 | 7 |
| *Quercus sadleriana* | 4 | 4 | 0 |
| *Quercus sagraeana* | 4 | 9 | 5 |
| *Quercus salicina* | 1 | 1 | 0 |
| *Quercus sapotifolia* | 2 | 2 | 0 |
| *Quercus sartorii* | 3 | 17 | 14 |
| *Quercus schottkyana* | 1 | 1 | 0 |
| *Quercus scytophylla* | 4 | 4 | 0 |
| *Quercus segoviensis* | 1 | 1 | 0 |
| *Quercus semecarpifolia* | 1 | 1 | 0 |
| *Quercus senescens* | 1 | 1 | 0 |
| *Quercus serrata* | 1 | 1 | 0 |
| *Quercus sessilifolia* | 1 | 1 | 0 |
| *Quercus setulosa* | 1 | 1 | 0 |
| *Quercus shumardii* | 3 | 3 | 0 |
| *Quercus sideroxyla* | 4 | 4 | 0 |
| *Quercus similis* | 1 | 1 | 0 |
| *Quercus sinuata* | 2 | 2 | 0 |
| *Quercus stellata* | 6 | 16 | 10 |
| *Quercus stewardiana* | 1 | 1 | 0 |
| *Quercus striatula* | 7 | 8 | 1 |
| *Quercus suber* | 3 | 3 | 0 |
| *Quercus subspathulata* | 3 | 3 | 0 |
| *Quercus texana* | 2 | 4 | 2 |
| *Quercus tomentella* | 2 | 2 | 0 |
| *Quercus toumeyi* | 2 | 2 | 0 |
| *Quercus trojana* | 1 | 1 | 0 |
| *Quercus turbinella* | 2 | 4 | 2 |
| *Quercus urbanii* | 2 | 4 | 2 |
| *Quercus utilis* | 1 | 1 | 0 |
| *Quercus uxoris* | 2 | 2 | 0 |
| *Quercus vacciniifolia* | 3 | 7 | 4 |
| *Quercus variabilis* | 2 | 2 | 0 |
| *Quercus vaseyana* | 2 | 2 | 0 |
| *Quercus velutina* | 2 | 2 | 0 |
| *Quercus viminea* | 2 | 2 | 0 |
| *Quercus virginiana* | 5 | 5 | 0 |
| *Quercus vulcanica* | 1 | 1 | 0 |
| *Quercus wislizeni* | 13 | 16 | 3 |
| *Quercus yiwuensis* | 1 | 1 | 0 |
| *Quercus yunnanensis* | 1 | 1 | 0 |

**Table S4** PhiIC values for alternative calibrations

| lambda | 0 | 0.05 | 0.1 | 0.5 | 1 | 5 | 10 |
| --- | --- | --- | --- | --- | --- | --- | --- |
| logLik | -5.12 | -5.12 | -5.14 | -5.53 | -6.24 | -8.24 | -7.77 |
| k | 755 | 755 | 755 | 755 | 755 | 755 | 755 |
| PHIIC-relaxed | 1520.24 | 1520.73 | 1521.25 | 1525.88 | 1532.09 | 1574.49 | 1621.70 |
| PHIIC-correlated | 1520.24 | 1520.24 | 1520.24 | 1520.24 | 1520.24 | 1520.24 | 1520.24 |

**Table S5** Phypart components and clade ages

| **Taxon** | **Age - stem calibration** | **Age - Crown calibration** | **Descendant** | **Concord** | **Conflict** | **% concord** |
| --- | --- | --- | --- | --- | --- | --- |
| 01. Quercus | 51.49 | 56.00 | 253 | 701 | 354 | 66.45% |
| 02. subg. *Quercus* | 47.87 | 54.14 | 180 | 129 | 640 | 16.78% |
| 03. subg. *Cerris* | 48.32 | 53.87 | 73 | 131 | 676 | 16.23% |
| 04. sect. *Quercus* | 28.11 | 45.00 | 92 | 108 | 482 | 18.31% |
| 05. sect. *Protobalanus* | 23.23 | 25.80 | 5 | 6 | 101 | 5.61% |
| 06. sect. *Virentes* | 22.25 | 31.45 | 7 | 108 | 49 | 68.79% |
| 07. Roburoid white oaks | 7.58 | 12.08 | 24 | 24 | 204 | 10.53% |
| 08. Mexican white oaks | 7.31 | 11.97 | 36 | 3 | 338 | 0.88% |
| 09. sect. Lobatae | 39.20 | 47.87 | 74 | 204 | 228 | 47.22% |
| 10. Laurifoliae | 17.72 | 19.94 | 16 | 8 | 234 | 3.31% |
| 11. Rubrae | 21.27 | 23.93 | 7 | 2 | 138 | 1.43% |
| 12. Mexican red oaks | 15.96 | 17.95 | 41 | 2 | 264 | 0.75% |
| 13. sect. *Cerris* | 21.87 | 37.82 | 14 | 261 | 144 | 64.44% |
| 14. sect. *Ilex* | 24.92 | 40.36 | 25 | 38 | 377 | 9.16% |
| 15. sect. *Cyclobalanopsis* | 36.24 | 48.32 | 34 | 134 | 321 | 29.45% |

**Methods S1** Analysis details: full methods for paper; abbreviated methods included in main text

### Previously published RAD-seq and new RAD-seq: sequencing and clustering

Data from several previously published RAD-seq phylogenies (Jiang *et al.*, In review; Cavender-Bares *et al.*, 2015; Hauser *et al.*, 2017; Fitz-Gibbon *et al.*, 2017; Pham *et al.*, 2017; McVay *et al.*, 2017a,a; Hipp *et al.*, 2018; Deng *et al.*, 2018) were analyzed alongside new RAD-seq data for a total of 795 sequencing runs for initial clustering analysis, reduced to 632 individuals after removal of technical replicates, poor sequencing reads, and individuals with insufficient vouchers and / or taxonomic data. RAD-seq data were generated as described in the previous studies. New data were from library preparations conducted at Floragenex, Inc. (Portland, OR, USA) following the methods of Baird et al. (2008) with *Pst*I, barcoded by individual, and sequenced on an Illumina Genome Analyzer IIx at Floragenex, or an Illumina HiSeq 2000, 2500 or 4000 at the University of Oregon Genomic Facility. Sequencing reads were 100 or 150 bp in length depending on the sequencing platform.

FASTQ files were demultiplexed without allowing any mismatches in the 10-bp barcode sequence and filtered to remove sequences with more than 5 bases of quality score < 20. Sequence data were assembled into loci for phylogenetic analysis using ipyrad 0.7.23 (Eaton, 2014) at 85% sequence similarity, requiring a minimum of 6 FASTQ sequences per individual per locus, and excluding loci with a cluster depth of > 10,000 sequences, more than 2 alleles, or more than 5 Ns or 8 heterozygotic positions in the consensus sequence. Consensus sequences for each individual for each locus were then clustered across individuals, retaining loci present in at least 4 individuals and possessing a maximum of 20 SNPs and 8 indels across individuals. Analysis up to this step took approximately 55 days on an Intel Xeon E5-1660 v3 with 8 cores, running at 3.00 GHz, and recovered a total of 222,850 RADseq loci, with number of loci per individual averaging 4582.9 +/- 1999.4 (sd), with a median of 4593 and a range of 26 to 10,765. More than half (113,244) of these loci were present in only 4 to 7 individuals. The dataset was winnowed down to loci with a minimum of 15 individuals each, for a total of 58,985 loci. Data were imported into R using the RADami package (Hipp *et al.*, 2014) for downstream analysis.

RAD-seq loci were mapped back to the latest version of the *Quercus robur* haploid genome (haplome 2.3; <https://urgi.versailles.inra.fr/Data/Genome/Genome-data-access>) (Plomion *et al.*, 2018). The oak genome is made of 12 pseudomolecules (*i.e*. chromosomes) and a set of 538 unassigned scaffolds. Mapping was performed using Blast+ 2.8.1 (Camacho *et al.*, 2009). We filtered alignments based on expect (E) values (E-value ≤10^-5^), alignment length (≥80% of the length of the loci) and percent identity (≥80%). For each locus, the best alignment was kept. Chromosome lengths, gene positions, and gene lengths were also taken from the latest version of the *Quercus robur* haploid genome as annotated (Plomion *et al.*, 2018, supplementary tables). Overlap of genes with mapped RAD-seq loci was quantified in R using the IRanges package (Lawrence *et al.*, 2013).

All sequence data analyzed in this paper are available as FASTQ files from NCBI’s Short Read Archive (Table S1), and aligned loci are available from https://github.com/andrew-hipp/global-oaks-2019.

### Phylogenetic analysis

Maximum likelihood phylogenetic analyses were conducted in RAxML v8.2.4 (Stamatakis, 2014) using the GTRCAT implementation of the general time reversible model of nucleotide evolution, with branch support assessed using the RELL bootstrapping (BS) option. GTRCAT applies the general-time reversible model of nucleotide evolution, with per-site variation modeled using the PSR (‘per-site rate’) approximation of the Gamma distribution (Stamatakis, 2006). Analyses were conducted on datasets clustered with a minimum of 15 individuals per locus; initial trials (not presented here) demonstrated that varying this number from 10 to 40 individuals per locus had little effect on the topology. For the phylogeny including all tips (Fig. S1), analysis was unconstrained, and topology within the white oaks of sections *Ponticae*, *Virentes*, and *Quercus* (hereafter in the paper “white oaks *s.l.*,” contrasted with “white oaks *s.s.*” for just section *Quercus*) was observed to be at odds with previous close studies (Crowl *et al.*, In review; McVay *et al.*, 2017b,a; Hipp *et al.*, 2018) that have shown the topology of the white oaks *s.l.* to be sensitive to taxon and locus sampling. For dating, samples were pruned to one sample per named species, favoring samples with the most loci samples, except for species in which variable position of samples from different populations is deemed to represent cryptic diversity, in which case more than one exemplar was retained. The singletons tree was estimated in RAxML using the following phylogenetic constraint:

(Ponticae, (Virentes, (Dumosae, (Stellatae + Mexicanae, (Prinoideae, (Albae, (Roburoids)))))))

Sectional nomenclature follows Denk et al. (2017) and the infrasectional names Manos (2016). The remainder of the tree was unconstrained and conforms closely to previous topologies.

We utilized neighbor-net (Bryant & Moulton, 2004) to visualize overall patterns of molecular genetic diversity. Likelihood-based methods that estimate species networks by explicitly modeling both lineage sorting and introgression as sources of among-locus incongruence (e.g., Solís-Lemus & Ané, 2016; Solís-Lemus *et al.*, 2017; Wen *et al.*, 2018; Zhang *et al.*, 2018) provide direct tests of introgression and hybridization hypotheses, allowing researchers to tease apart the relative contributions to gene tree incongruence of stochastic allele sorting on one hand, allele migration among lineages on the other. However, such approaches are not currently tractable with large numbers of taxa analyzed simultaneous, and those that start with gene trees lose power with short-read RAD-seq data. We have utilized such methods on smaller numbers of taxa within the oaks (Crowl *et al.*, In review; Eaton *et al.*, 2015; Hauser *et al.*, 2017; McVay *et al.*, 2017b,a), but for the current study we utilize a simple distance-based approach that yields a planar (2-dimensional) splits network as implemented in SplitsTree v. 14.3 (Huson & Bryant, 2006). The neighbor-net, as used here, can be seen as a meta-phylogenetic network visualizing in two dimensions the approximate range of (semi-compatible) trees that are implied under a distance criterion by our RAD-seq dataset. The splits network was inferred from a maximum-likelihood (GTR+Γ) pairwise distance matrix estimated in RAxML, using the same datasets utilized for the singletons tree, excluding the non-*Quercus* outgroups.

### Calibration of singletons tree

Branch lengths on the tree were inferred using penalized likelihood under both a relaxed model, where rates are uncorrelated among branches (Paradis, 2013); and a correlated rates model (which corresponds to the penalized likelihood approach of Sanderson, 2002), as implemented in the chronos function of ape v 5.1 (Paradis *et al.*, 2004) of R v 3.4.4 (“Someone to Lean On”) (R-Development-Core-Team, 2004). Nodes were calibrated in two different ways, either using eight fossil calibrations, corresponding to the crown of the genus, the crown of two sections in subgenus *Quercus* and two in subgenus *Cerris*, one calibration within section *Ilex* and two within *Cerris* (Table S2) (McIntyre, D.J., 1991; Manchester, 1994, 2011; Tanai & Uemura, 1994; McIver & Basinger, 1999; Akhmetiev *et al.*, 2009; Hofmann, 2010; Standke *et al.*, 2010; Kmenta, 2011; Hofmann *et al.*, 2011; Denk *et al.*, 2012; Eberle & Greenwood, 2012; Bouchal *et al.*, 2014; Pavlyutkin, 2015; Greenwood *et al.*, 2016; Grímsson *et al.*, 2016; Su *et al.*, 2018); or more conservatively as stem ages, using a subset of five fossils corresponding to the stem of the genus, of sections *Lobatae*, *Cyclobalanopsis*, and *Quercus*, and section *Cerris* in part. Fossils were assigned to clades based on possession of morphological synapomorphies. The two alternative sets of calibrations account for uncertainty in our knowledge of how fully the fossil record is known (i.e., there may be older fossils that we missed, justifying a crown calibration) and in the placement along the stem of each fossil. The former (hereafter referred to as the crown calibration) may slightly overestimate divergence ages but takes into account that, in reality, early fossils of a lineage are often closer to the most recent common ancestor of a group than to the earliest ancestor of the group (Forest, 2009). For all fossil priors except for the root of section *Quercus*, the difference in calibration points is one node. Thus our two calibrations bracket what we consider to be plausible age ranges for the tree, conditioned on placement of these fossils somewhere along the branch where they are placed (Table S2). Ages were treated as fixed (for precisely dated fossils) or with maximum and minimum of corresponding epochs / ages (Table S2). A separate estimate of the best fit λ for the correlated clock model was made using cross-validation as implemented in the chronopl function of ape, and that value of λ was used for both the relaxed and correlated clocks. Branch lengths were optimized over smoothing parameter (λ) values of 0, 0.05, 0.1, 0.5, 1, 5 and 10 to assess sensitivity of branch length inferences to balance between the parametric and non-parametric components of each model. Comparison of *ϕ*IC was used to identify the best fit model for each value of λ (Table S4).

Transitions in lineage diversification rates were estimated using the speciation-extinction model implemented in Bayesian Analysis of Macroevolutionary Mixtures (BAMM) (Rabosky, 2014); the BAMMtools R package was used for configuration and analysis of MCMC. Priors were set using the setBAMMpriors function. Four BAMM analyses were conducted: one each on the crown calibrations tree and the stem calibrations tree assuming a global sampling proportion of 60%; and one each on these same trees assuming separate sampling proportions by clade. The latter analyses account for the fact that while we have sampled an estimated 60% of the species in the genus overall, we have undersampled the Mexican / Central American / southwestern U.S. white oaks (ca. 51%) and red oaks (ca. 57%) and the Eurasian sections *Ilex* (69%) and *Cyclobalanopsis* (38%). Analyses were run for 4*10^6^ generations, saving every 2000 generations, with four chains per MCMC analysis. ESS for final ln L post-burnin was >1200, and several replicate runs yielded the same configuration of rate shifts, suggesting sufficient mixing. To visualize changes in standing diversity over time for the different sections, we plotted lineage through time (LTT) plots by section against δ^18^O levels reported in Zachos *et al.* (2001) as a proxy for mean global temperatures. While these levels can be converted to degrees centigrade (e.g., following the formula of Epstein *et al.* (1953), we leave them as unconverted δ^18^O values to avoid over-interpretation, given the high uncertainty in inferences of past surface climates from any temperature proxies (Lunt *et al.*, 2012).

### Investigating the genomic landscape of oak evolutionary history

Introgressive status of loci for two known introgression events involving the Eurasian white oaks (McVay *et al.*, 2017b) and the western North American lobed-leaf white oaks (McVay *et al.*, 2017a) was assessed by calculating the likelihood of phylogenies inferred for each locus under the constraint of the inferred divergence history (species tree) and the gene flow history at odds with that divergence history, as inferred in the studies cited above. These two cases are of particular interest because they are well studied, and lineage sorting has been ruled out in the above studies as an explanation of incongruence between the alternative topologies we test. Thus mapping this back to the genome gives a map of alternative locus histories that are a consequence of introgression. Likelihood ratios (log-likelihood difference) were used to assess the relative support of each locus for the introgression vs. the divergence history. Position of loci with a relative support of at least 2 log-likelihood points for one history relative to the other were mapped back to the *Quercus robur* genome as assembled into 12 pseudomolecules or “pseudochromosomes” (Plomion *et al.*, 2018). Loci were screened for this from the all-tips tree to maximize the number of loci that could be evaluated.

To identify relative phylogenetic informativeness of loci, two tests were conducted based on the singletons tree. First, the ML topology was estimated in RAxML for each of 2844 loci that were (1) mapped back to the *Quercus robur* genome, (2) sequenced for a minimum of 10 individuals in the singletons tree, and (3) rootable because they had at least one non-*Quercus* sample. Trees were read into R, and branches less than 10^-5^ changes / nucleotide were collapsed to make polytomies. At this point, trees with only one node (the root node) were discarded, leaving 2762 locus trees that resolved an average of 4.48 (+/- 1.83 s.d.) nodes each, with a maximum of 15 and a median of 4. These were then compared with the total-evidence tree using quartet similarities calculated under the tqDist algorithm (Sand *et al.*, 2014), using the Quartet package (Smith, 2019). We used as our similarity metric the number of quartets resolved the same way for both the locus tree and the whole singletons tree divided by the sum of quartets resolved the same and those resolved differently. This similarity metric ignores differences in taxon sampling between the locus trees and the whole tree and quartets that cannot be resolved in the locus trees. Quartet similarities were mapped back to the genome to identify regions of particularly high or low phylogenetic fidelity.

Second, these same locus trees were mapped back to the singletons tree using phyparts (Smith *et al.*, 2015), which identifies for all branches on a single tree how many individual locus trees support or reject that branch. We use these data to investigate the genomic pattern of incongruence between individual loci and the total evidence tree and to test the hypothesis that there is genomic autocorrelation in phylogenetic signal, which would indicate conserved regions of the genome that are important in divergence across the oak phylogeny. We test this globally, using spline correlograms (Bjørnstad & Falck, 2001; Bjørnstad, 2008) to estimate the spatial scale of nonindependence among congruence and incongruence for the tree as a whole, with each chromosome tested independently.

**REFERENCES CITED: Methods S1 and Table S2**

**Akhmetiev M, Walther H, Kvaček Z**. **2009**. Mid-Latitude Palaeogene Floras of Eurasia Bound to Volcanic Settings and Palaeoclimatic Events -- Experience Obtained from the Far East of Russia (sikhote-Alin’) and Central Europe (Bohemian Massif). *Acta Musei Nationalis Pragae, Series B - Historia Naturalis* **65**: 61–129.

**Baird NA, Etter PD, Atwood TS, Currey MC, Shiver AL, Lewis ZA, Selker EU, Cresko WA, Johnson EA**. **2008**. Rapid SNP Discovery and Genetic Mapping Using Sequenced RAD Markers. *PLOS ONE* **3**: e3376.

**Bjørnstad ON**. **2008**. *ncf: spatial nonparametric covariance functions*.

**Bjørnstad ON, Falck W**. **2001**. Nonparametric spatial covariance functions: Estimation and testing. *Environmental and Ecological Statistics* **8**: 53–70.

**Bouchal J, Zetter R, Grímsson F, Denk T**. **2014**. Evolutionary trends and ecological differentiation in early Cenozoic Fagaceae of western North America. *American Journal of Botany* **101**: 1332–1349.

**Bryant D, Moulton V**. **2004**. Neighbor-Net: An Agglomerative Method for the Construction of Phylogenetic Networks. *Molecular Biology and Evolution* **21**: 255–265.

**Camacho C, Coulouris G, Avagyan V, Ma N, Papadopoulos J, Bealer K, Madden TL**. **2009**. BLAST+: architecture and applications. *BMC Bioinformatics* **10**: 421.

**Cavender-Bares J, Gonzalez-Rodriguez A, Eaton DAR, Hipp AL, Beulke A, Manos PS**. **2015**. Phylogeny and biogeography of the American live oaks (*Quercus* subsection *Virentes*): A genomic and population genetics approach. *Molecular Ecology* **24**: 3668–3687.

**Crowl AA, McVay JD, Manos PS, Hipp AL, Lemmon A, Lemmon E**. **In review**. Uncovering the genomic signature of ancient introgression between white oak lineages (*Quercus*) using anchored enrichment. *New Phytologist*.

**Deng M, Jiang X-L, Hipp AL, Manos PS, Hahn M**. **2018**. Phylogeny and biogeography of East Asian evergreen oaks (*Quercus* section *Cyclobalanopsis*; Fagaceae): Insights into the Cenozoic history of evergreen broad-leaved forests in subtropical Asia. *Molecular Phylogenetics and Evolution* **119**: 170–181.

**Denk T, Grimm GW, Manos PS, Deng M, Hipp AL**. **2017**. An Updated Infrageneric Classification of the Oaks: Review of Previous Taxonomic Schemes and Synthesis of Evolutionary Patterns. In: Tree Physiology. Oaks Physiological Ecology. Exploring the Functional Diversity of Genus *Quercus* L. Springer, Cham, 13–38.

**Denk T, Grímsson F, Zetter R**. **2012**. Fagaceae from the early Oligocene of Central Europe: Persisting new world and emerging old world biogeographic links. *Review of Palaeobotany and Palynology* **169**: 7–20.

**Eaton DAR**. **2014**. PyRAD: assembly of de novo RADseq loci for phylogenetic analyses. *Bioinformatics (Oxford, England)* **30**: 1844–1849.

**Eaton DAR, Hipp AL, González-Rodríguez A, Cavender-Bares J**. **2015**. Historical introgression among the American live oaks and the comparative nature of tests for introgression. *Evolution* **69**: 2587–2601.

**Eberle JJ, Greenwood DR**. **2012**. Life at the top of the greenhouse Eocene world—A review of the Eocene flora and vertebrate fauna from Canada’s High Arctic. *GSA Bulletin* **124**: 3–23.

**Epstein S, Buchsbaum R, Lowenstam HA, Urey HC**. **1953**. Revised carbonate-water isotopic temperature scale. *GSA Bulletin* **64**: 1315–1326.

**Fitz-Gibbon S, Hipp AL, Pham KK, Manos PS, Sork V**. **2017**. Phylogenomic inferences from reference-mapped and de novo assembled short-read sequence data using RADseq sequencing of California white oaks (*Quercus* subgenus *Quercus*). *Genome* **60**: 743–755.

**Forest F**. **2009**. Calibrating the Tree of Life: fossils, molecules and evolutionary timescales. *Annals of Botany* **104**: 789–794.

**Greenwood DR, Pigg KB, Basinger JF, DeVore ML**. **2016**. A review of paleobotanical studies of the Early Eocene Okanagan (Okanogan) Highlands floras of British Columbia, Canada, and Washington, USA. *Canadian Journal of Earth Sciences* **53**: 548–564.

**Grímsson F, Grimm GW, Zetter R, Denk T**. **2016**. Cretaceous and Paleogene Fagaceae from North America and Greenland: evidence for a Late Cretaceous split between Fagus and the remaining Fagaceae. *Acta Palaeobotanica* **56**: 247–305.

**Hauser DA, Keuter A, McVay JD, HIpp AL, Manos PS**. **2017**. The evolution and diversification of the red oaks of the California Floristic Province (*Quercus* section *Lobatae*, series *Agrifoliae*). *American Journal of Botany* **104**: 1581–1595.

**Hipp AL, Eaton DAR, Cavender-Bares J, Fitzek E, Nipper R, Manos PS**. **2014**. A framework phylogeny of the American oak clade based on sequenced RAD data. *PLoS ONE* **9**: e93975.

**Hipp AL, Manos PS, González-Rodríguez A, Hahn M, Kaproth M, McVay JD, Avalos SV, Cavender-Bares J**. **2018**. Sympatric parallel diversification of major oak clades in the Americas and the origins of Mexican species diversity. *New Phytologist* **217**: 439–452.

**Hofmann C-C**. **2010**. Microstructure of Fagaceae pollen from Austria (Paleocene/Eocene boundary) and Hainan Island (middle Eocene). In: 8th European Palaeobotany-Palynology Conference. Budapest: Hungarian Natural History Museum, 119.

**Hofmann C-C, Mohamed O, Egger H**. **2011**. A new terrestrial palynoflora from the Palaeocene/Eocene boundary in the northwestern Tethyan realm (St. Pankraz, Austria). *Review of Palaeobotany and Palynology* **166**: 295–310.

**Huson DH, Bryant D**. **2006**. Application of Phylogenetic Networks in Evolutionary Studies. *Molecular Biology and Evolution* **23**: 254–267.

**Jiang X-L, Deng M, Hipp AL, Su T, Yan M-X, Zho Z-K**. **In review**. East Asian origins of European holly oaks via the Tibet-Himalayas. *Journal of Biogeography*.

**Kmenta M**. **2011**. Die Mikroflora der untermiozänen Fundstelle Altmittweida, Deutschland.

**Lawrence M, Huber W, Pagès H, Aboyoun P, Carlson M, Gentleman R, Morgan MT, Carey VJ**. **2013**. Software for Computing and Annotating Genomic Ranges. *PLOS Computational Biology* **9**: e1003118.

**Lunt DJ, Dunkley Jones T, Heinemann M, Huber M, LeGrande A, Winguth A, Loptson C, Marotzke J, Roberts CD, Tindall J, *et al.*** **2012**. A model–data comparison for a multi-model ensemble of early Eocene atmosphere–ocean simulations: EoMIP. *Climate of the Past* **8**: 1717–1736.

**Manchester SR**. **1994**. Fruits and Seeds of the Middle Eocene Nut Beds Flora, Clarno Formation, Oregon. *Palaeontographica Americana* **58**: 1–205.

**Manchester SR**. **2011**. Fruits of Ticodendraceae (Fagales) from the Eocene of Europe and North America. *International Journal of Plant Sciences* **172**: 1179–1187.

**Manos PS**. **2016**. Systematics and biogeography of the American oaks. *International Oaks* **27**: 23–36.

**McIntyre, D.J.** **1991**. Pollen and spore flora of an Eocene forest, eastern Axel Heiberg Island. *N.W.T. Geological Survey of Canada Bulletin* **403**: 83–97.

**McIver EE, Basinger JF**. **1999**. Early Tertiary Floral Evolution in the Canadian High Arctic. *Annals of the Missouri Botanical Garden* **86**: 523–545.

**McVay JD, Hauser D, Hipp AL, Manos PS**. **2017a**. Phylogenomics reveals a complex evolutionary history of lobed-leaf white oaks in western North America. *Genome* **60**: 733–742.

**McVay JD, Hipp AL, Manos PS**. **2017b**. A genetic legacy of introgression confounds phylogeny and biogeography in oaks. *Proc. R. Soc. B* **284**: 20170300.

**Paradis E**. **2013**. Molecular dating of phylogenies by likelihood methods: A comparison of models and a new information criterion. *Molecular Phylogenetics and Evolution* **67**: 436–444.

**Paradis E, Claude J, Strimmer K**. **2004**. APE: Analyses of Phylogenetics and Evolution in R language. *Bioinformatics* **20**: 289–290.

**Pavlyutkin BI**. **2015**. The genus *Quercus* (Fagaceae) in the Early Oligocene Flora of Kraskino, Primorskii Region. *Paleontological Journal* **49**: 668–676.

**Spicer RA, Herman AB, Liao W, Spicer TEV, Kodrul TM, Yang J, Jin J**. **2014**. Cool tropics in the Middle Eocene: Evidence from the Changchang Flora, Hainan Island, China. *Palaeogeography, Palaeoclimatology, Palaeoecology* **412**: 1–16.

**Stamatakis A**. **2014**. RAxML Version 8: A tool for Phylogenetic Analysis and Post-Analysis of Large Phylogenies. *Bioinformatics* **30**: 1312–1313.

**Standke G, Escher D, Fischer J, Rascher J**. **2010**. *Das Tertiär Nordwestsachsens: Ein geologischer Überblick*. Freiberg: Sächsisches Landesamt für Umwelt, Landwirtschaft und Geologie.

**Su T, Spicer RA, Li S-H, Xu H, Huang J, Sherlock S, Huang Y-J, Li S-F, Wang L, Jia L-B, *et al.*** **2018**. Uplift, climate and biotic changes at the Eocene–Oligocene transition in south-eastern Tibet. *National Science Review*.

**Tanai T, Uemura K**. **1994**. Lobed oak leaves from the Tertiary of East Asia with reference to the oak phytogeography of the northern hemisphere. *Transactions and proceedings of the Paleontological Society of Japan. New series* **1994**: 343–365.

**Wen D, Yu Y, Zhu J, Nakhleh L**. **2018**. Inferring Phylogenetic Networks Using PhyloNet. *Systematic Biology* **67**: 735–740.

**Zachos J, Pagani M, Sloan L, Thomas E, Billups K**. **2001**. Trends, Rhythms, and Aberrations in Global Climate 65 Ma to Present. *Science* **292**: 686–693.

**Zhang C, Ogilvie HA, Drummond AJ, Stadler T**. **2018**. Bayesian Inference of Species Networks from Multilocus Sequence Data. *Molecular Biology and Evolution* **35**: 504–517.
