## Supplementary figures and images for "Genomic landscape of the global oak phylogeny"

### Fig S1 -- all-tips phylogeny, multiple pages

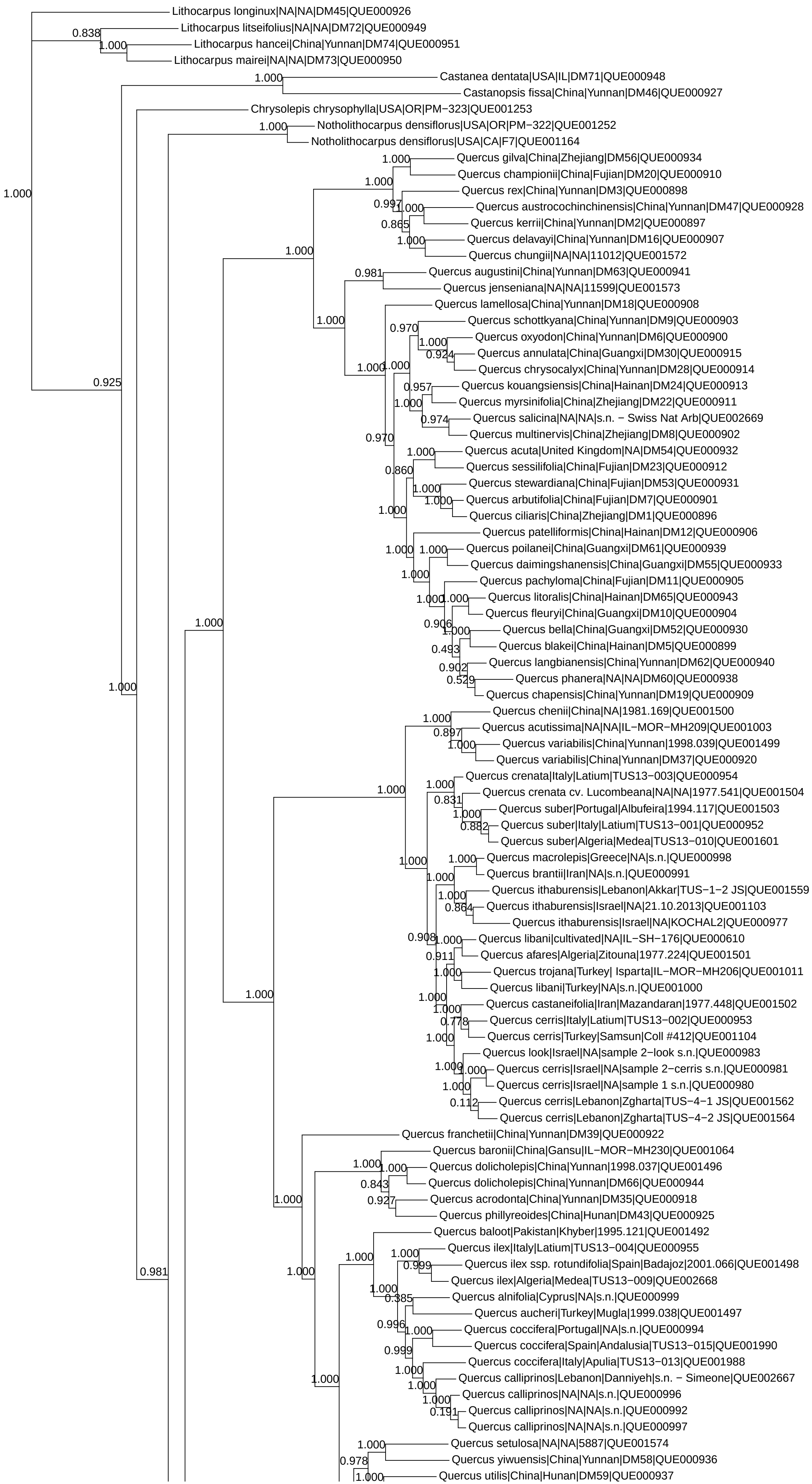

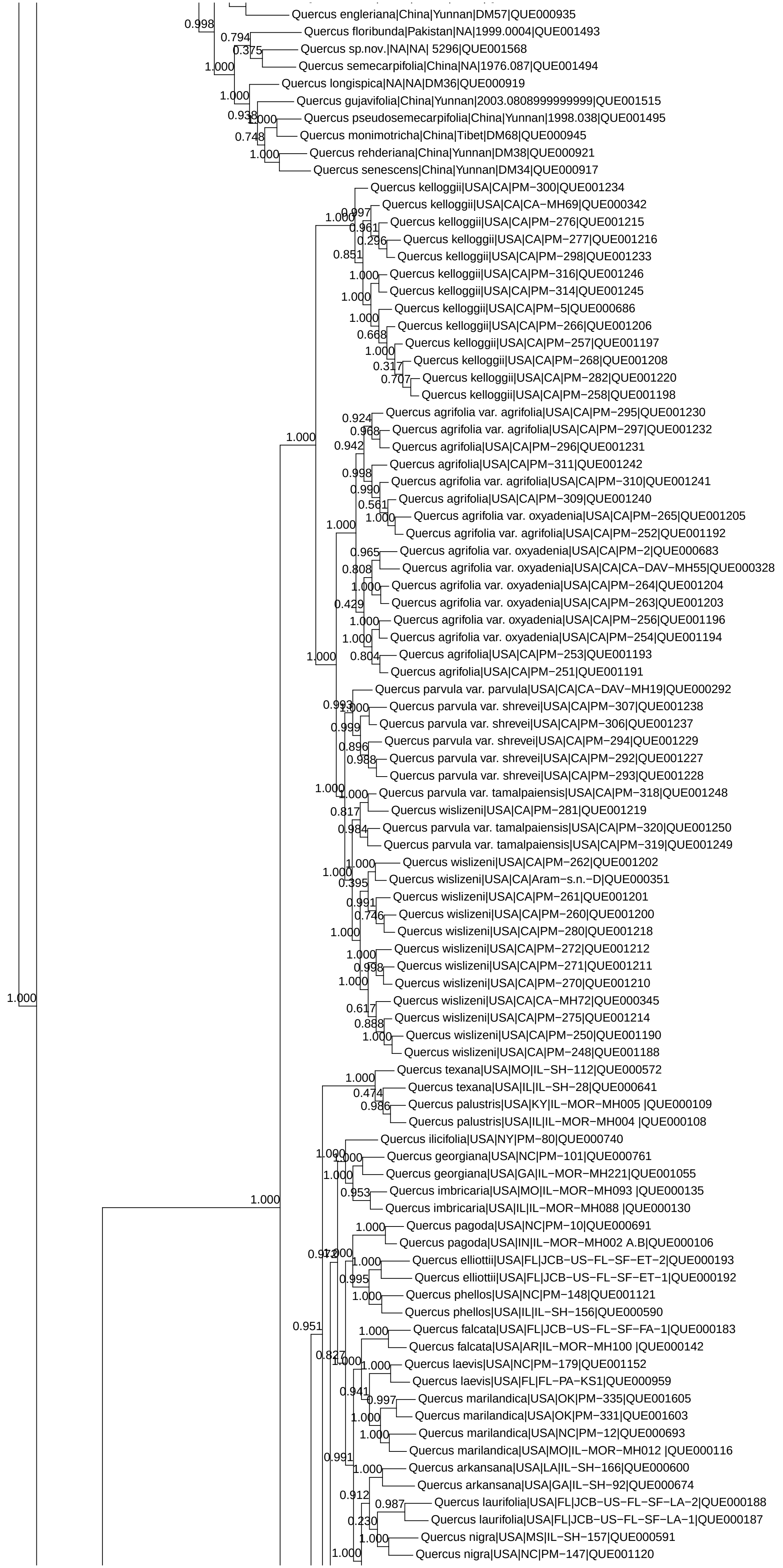

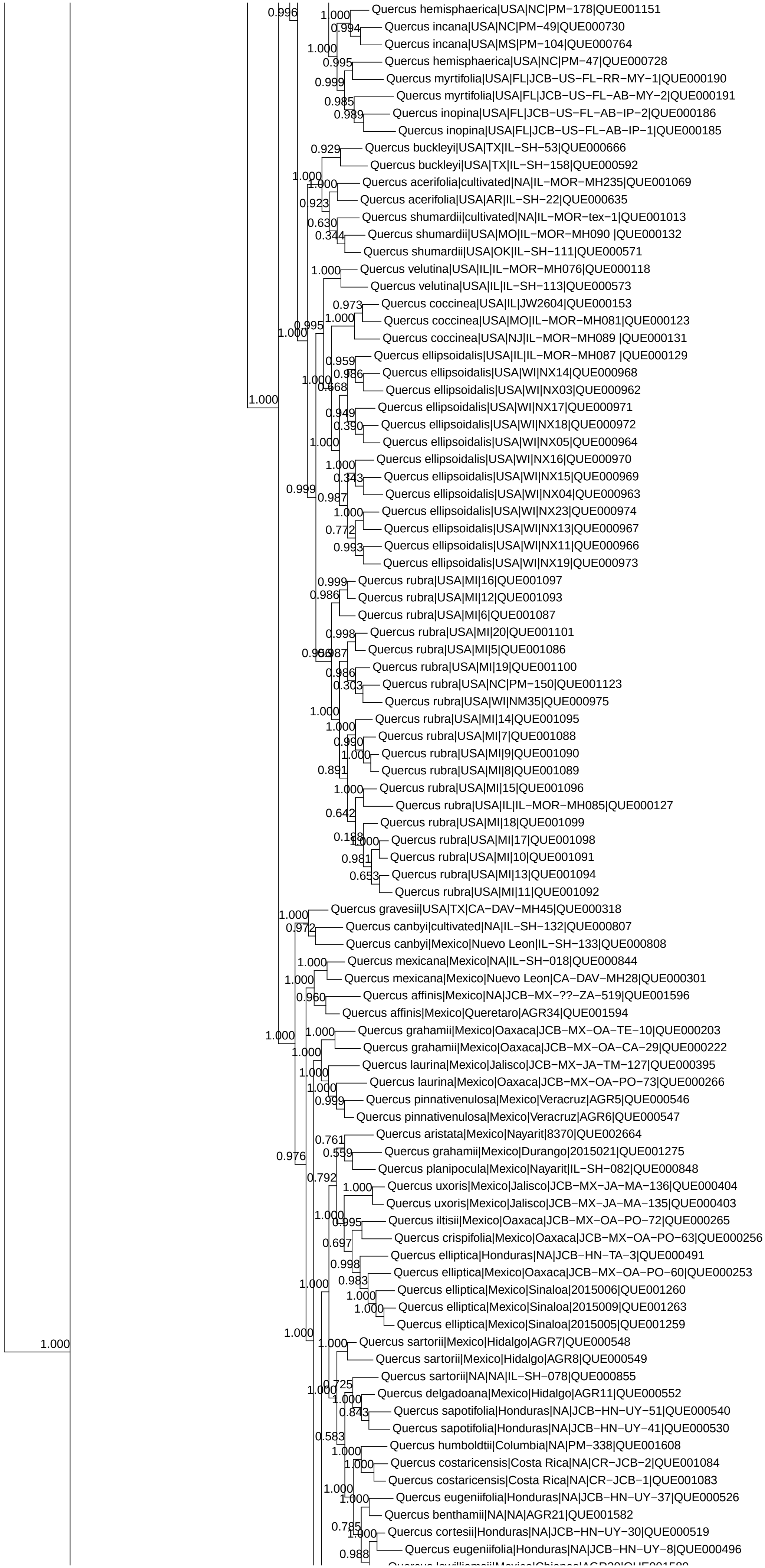

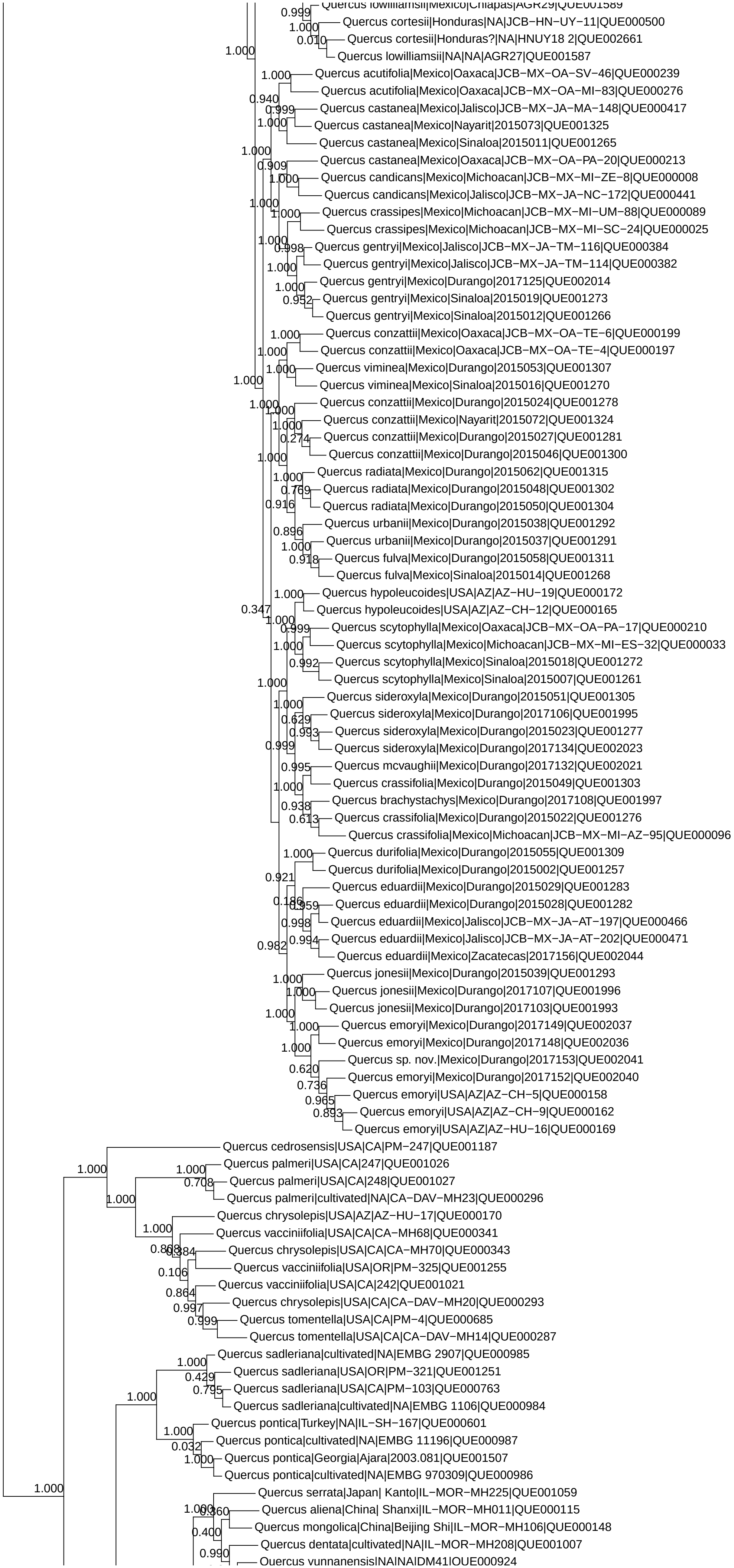

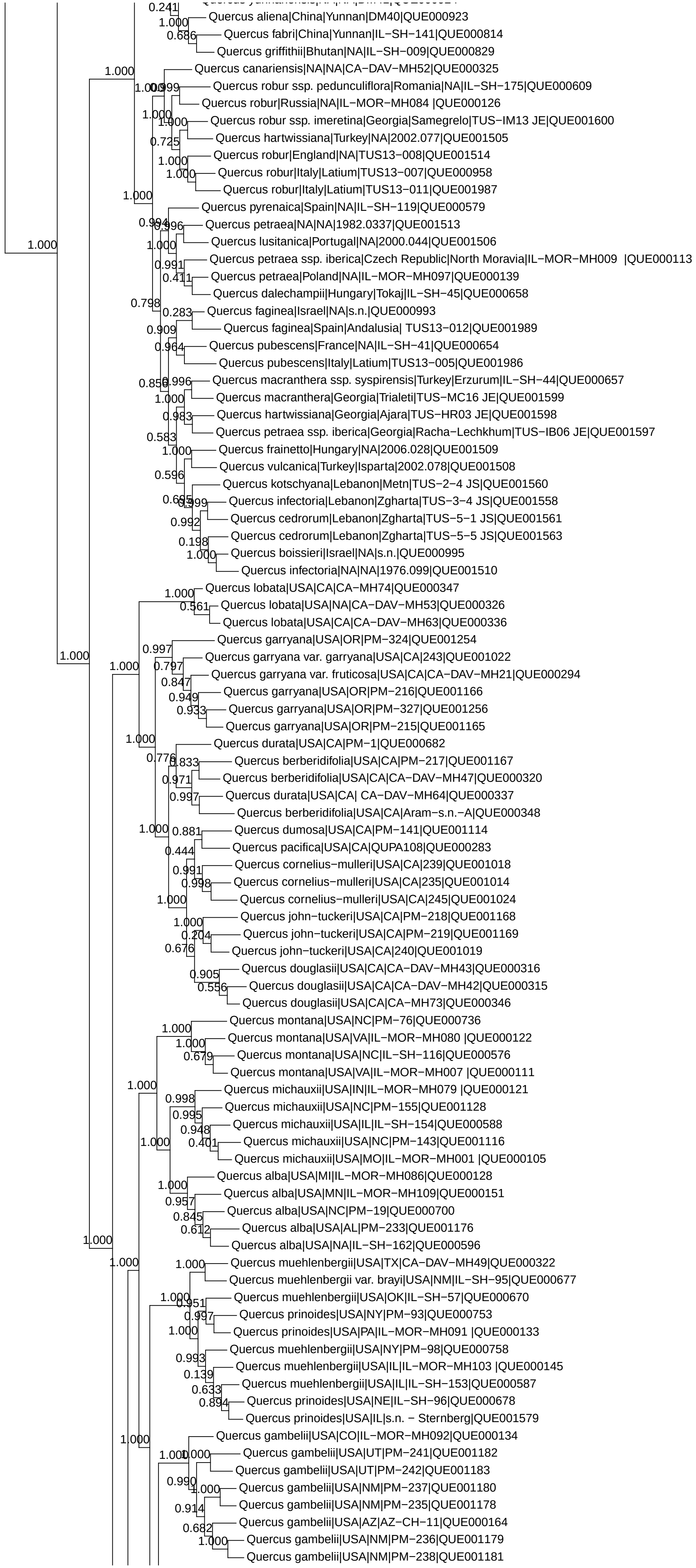

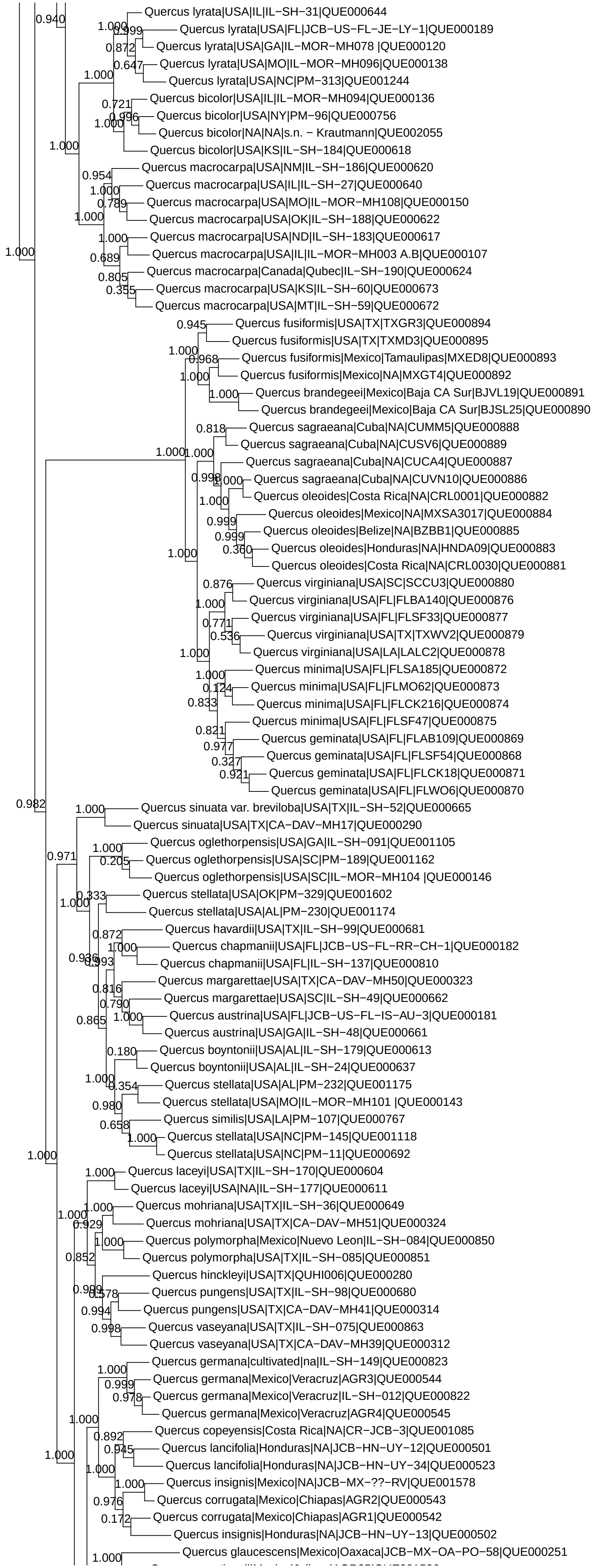

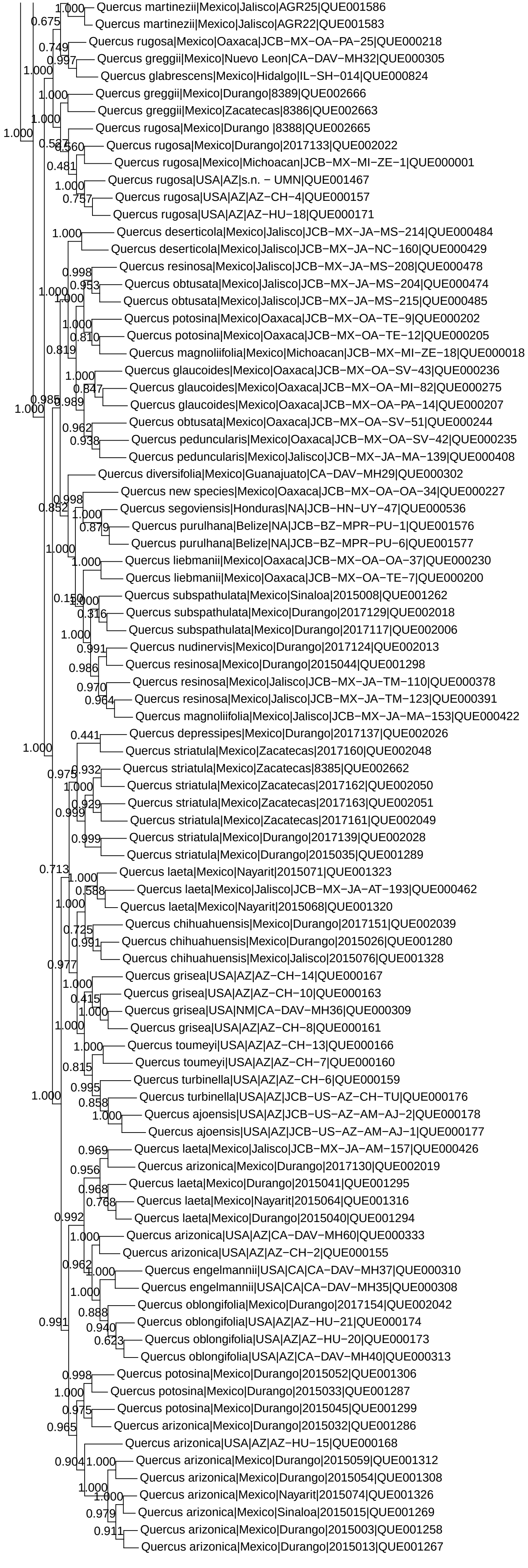
